# Bclaf1 biomolecular condensates protect nuclear PTK2 from ubiquitin-proteasome system promoting cardiomyocyte survival during oxidative stress

**DOI:** 10.1101/2025.02.04.636487

**Authors:** Isabela Aparecida Moretto, Beatriz Rocha Ilidio Rodrigues, Pedro Víctor-Carvalho, Maria das Graças de Souza Carvalho, Mariana Conceição da Silva, Fernando Valdivieso-Rivera, Giovanna Lopes de Araújo, Ana Paula Samogim, Lara Basseres Novais, Ingridi Rafaela de Brito, Guilherme Reis-de-Oliveira, Alan Gonçalves Amaral, Mariana Ozello Baratti, Fernanda Luisa Basei, Murilo Vieira Geraldo, Paulo Costa Carvalho, Marlon Dias Mariano Santos, Rosario Duran, Carlos Henrique Grossi Sponton, Jörg Kobarg, Fabio C. Gozzo, Hernandes F. de Carvalho, Andre Alexandre de Thomaz, Aline Mara dos Santos

## Abstract

PTK2, a non-receptor tyrosine kinase, plays a critical role in regulating essential cellular functions, including cell survival, by reducing p53 levels and activating the PI3K/AKT pathway. However, the mechanism underlying PTK2 stabilization during stress remains unclear. In this study, we identified Bclaf1, a multifunctional protein known to stabilize partners, as a PTK2 interactor. Using advanced microscopy techniques we identified nuclear Bclaf1 biomolecular condensates containing PTK2 in cardiomyocytes under oxidative stress. While diffuse PTK2 in the nucleus was susceptible to ubiquitination, PTK2 sequestered in the Bclaf1 condensates was protected from the ubiquitin-proteasome system (UPS). The K926 residue was identified as a ubiquitination site on PTK2, and subsequent proteasome inhibition experiments confirmed the role of the UPS in PTK2 homeostasis. Furthermore, disrupting Bclaf1 biomolecular condensates lead to PTK2 degradation, subsequently increasing p53 levels and activating apoptosis. Our findings support the role of Bclaf1 in the formation of pro-survival nuclear condensates that sequester and stabilize PTK2, promoting cardiomyocyte survival during oxidative stress.

**Graphical Abstract:** 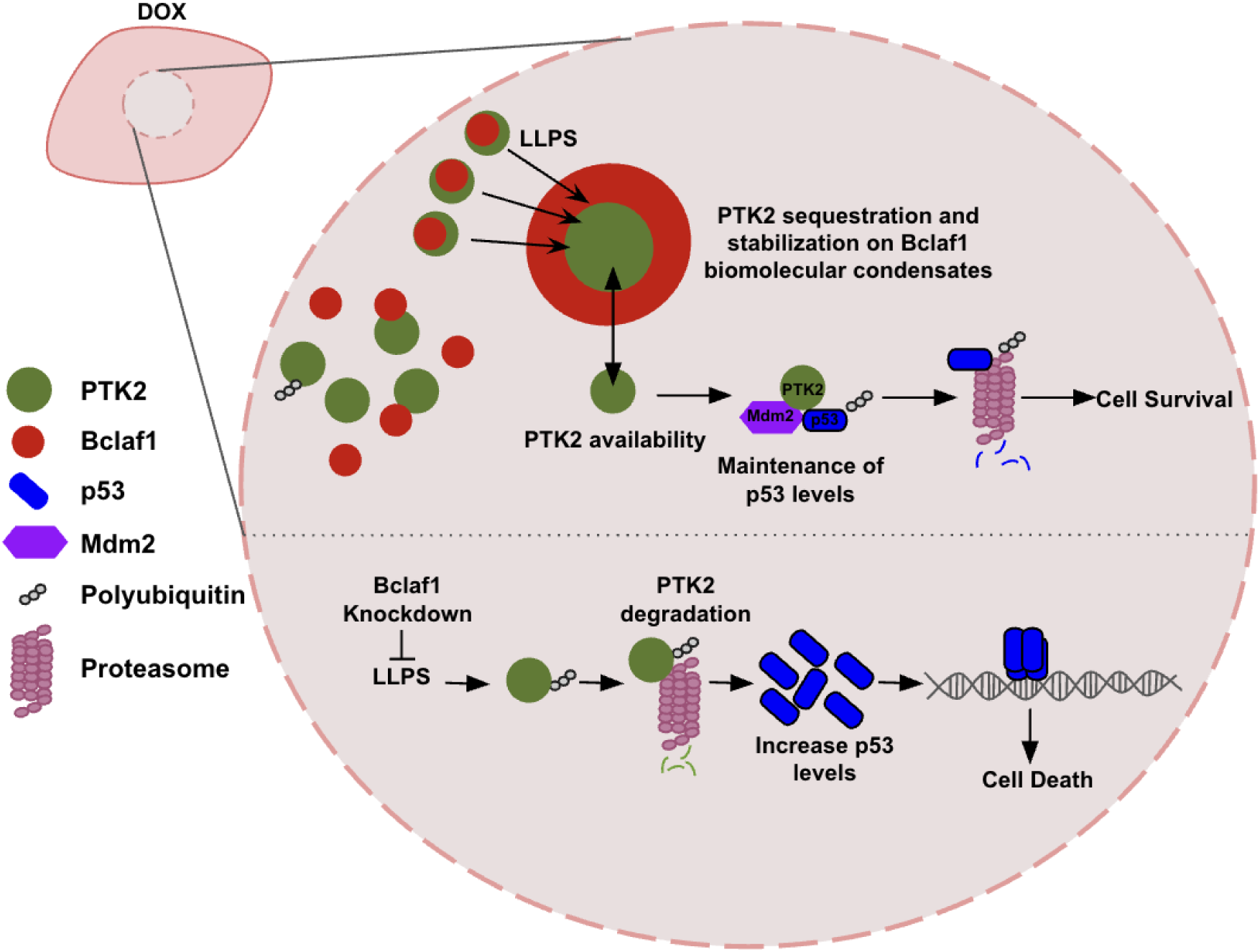

**In Brief:** Moretto et al. demonstrated that Bclaf1 undergo liquid-liquid phase separation (LLPS) during doxorubicin (Dox) induced oxidative stress to generate membraneless organelles where PTK2 is sequestered and protected from the ubiquitin-proteasome system (UPS). The maintenance of PTK2 stability may favor the p53 and Mdm2 interaction, which results in p53 ubiquitination and degradation, promoting cell survival. Loss of Bclaf1 disrupts PTK2 stabilization, leading to the ubiquitination and degradation of this kinase and downstream upregulation of p53, increasing cell death.

**Highlights:**

- Bclaf1 undergoes LLPS to stabilize protein partners during stress.
- PTK2 inside the Bclaf1 biomolecular condensates is protected from the UPS to promote cardiomyocyte survival during dox-induced oxidative stress.
- The PTK2 ubiquitination site, lysine 926, was identified and the action of the UPS on PTK2 proteostasis was confirmed.
- Bclaf1 knockdown results in overall protein ubiquitination and in PTK2 ubiquitination and degradation, resulting in increased levels of p53 and PUMA.

## 1. INTRODUCTION

Protein Tyrosine Kinase 2 (PTK2), also known as Focal Adhesion Kinase (FAK), is a multifunctional non-receptor tyrosine kinase involved in the regulation of primary cellular functions, including migration, mechanosensing, proliferation, and survival^1^. Under stress conditions, including mechanical and oxidative stress, PTK2 is rapidly activated through tyrosine 397 autophosphorylation^2,3,4^. Once phosphorylated, PTK2 recruits Src kinases to phosphorylate PTK2 on tyrosines 576 and 577 in the kinase activation loop, becoming fully activated^5,75,7^. Active PTK2 redistributes throughout the cytoplasm and nucleus to regulate several signaling pathways, such as PI3K-AKT-mTOR, Mdm2-p53, and MEF2c-cJun, through its kinase activity or scaffolding function, promoting cell survival and hypertrophic growth of cardiomyocytes^2,8–12^.

Several reports indicate nuclear PTK2 plays pivotal roles in the regulation of cell survival under stress^13–15^. Specific cardiomyocyte PTK2-knockout mice showed increased susceptibility to apoptosis and cardiac dysfunction after doxorubicin (dox)-induced oxidative stress, while the myocyte specific overexpression of PTK2 enhanced cell survival, providing cardioprotection to dox treatment^13^. With oxidative stress generated by dox and the key role of PTK2 to promote cardioprotection in mind, we reasoned that PTK2 must be protected and stabilized in cardiomyocytes to promote survival during oxidative stress conditions. By an unbiased immunoprecipitation coupled to mass spectrometry approach, we identified the Bcl-2-associated transcription factor 1 (Bclaf1) as one of the most enriched interacting partners of the FERM (four-point-one, ezrin, radixin, moesin)-PTK2 domain. Bclaf1 was previously shown to function on the stabilization and proteostasis control of interacting-partners under stress conditions^16,17^. However, the mechanism of protein stabilization by Bclaf1 remained uncharacterized.

In this study, using a combination of biochemical and biophysical assays with super-resolution microscopy, we identified nuclear Bclaf1 biomolecular condensates sequestering and protecting PTK2 from the ubiquitin-proteasome system during doxorubicin-induced oxidative stress, into submicron-scale membraneless organelles. Fluorescence Recovery After Photobleaching (FRAP) revealed the ability of Bclaf1 to form biomolecular condensates through liquid–liquid phase separation (LLPS). Knockdown of Bclaf1 resulted in increased PTK2 ubiquitination and degradation, and in cell death through activation of apoptosis pathways. Moreover, a PTK2 lysine ubiquitination site was identified by mass spectrometry, while proteasome inhibition resulted in nuclear PTK2 accumulation and ubiquitination, confirming the participation of the ubiquitin-proteasome system (UPS) on PTK2 homeostasis. Here, we present a new PTK2 quality-control mechanism that operates during oxidative stress to stabilize PTK2 and promote cell survival. Such mechanisms may promote cardioprotection during stress conditions, including dox-induced cytotoxicity through oxidative stress.

## 2. RESULTS

### 2.1 Bclaf1 is a novel nuclear binding partner of PTK2

Protein-protein interactions regulate a wide range of cell functions during physiological or pathological conditions, with its native structural conformation being essential to their proper functionality^18,19^. In order to identify PTK2 stabilizing interacting partners, we conducted an immunoprecipitation (IP) coupled to mass spectrometry. As a result, Bclaf1 was one of the most enriched interacting partners of the FERM-PTK2 domain in HEK-293T cells (Fig. 1A; Supplem. Table 1). Interactome data bank analysis showed PTK2 and Bclaf1 linked to overlapping pathways involved in cell signaling, however, this is the first time this interaction was described (Fig. 1B; Supplem. Fig. 1). Immunoprecipitation of PTK2 from lysates of H9c2 cardiomyocytes confirmed the PTK2-Bclaf1 association in this cell type (Fig. 1C; Supplem. Fig. 7A). When Bclaf1 was immunoprecipitated, PTK2 also was found to co-precipitate (Fig 1D; Supplem. Fig. 7A). This validates the PTK2-Bclaf1 interaction in H9c2 cardiomyocytes.

**Figure 1.**
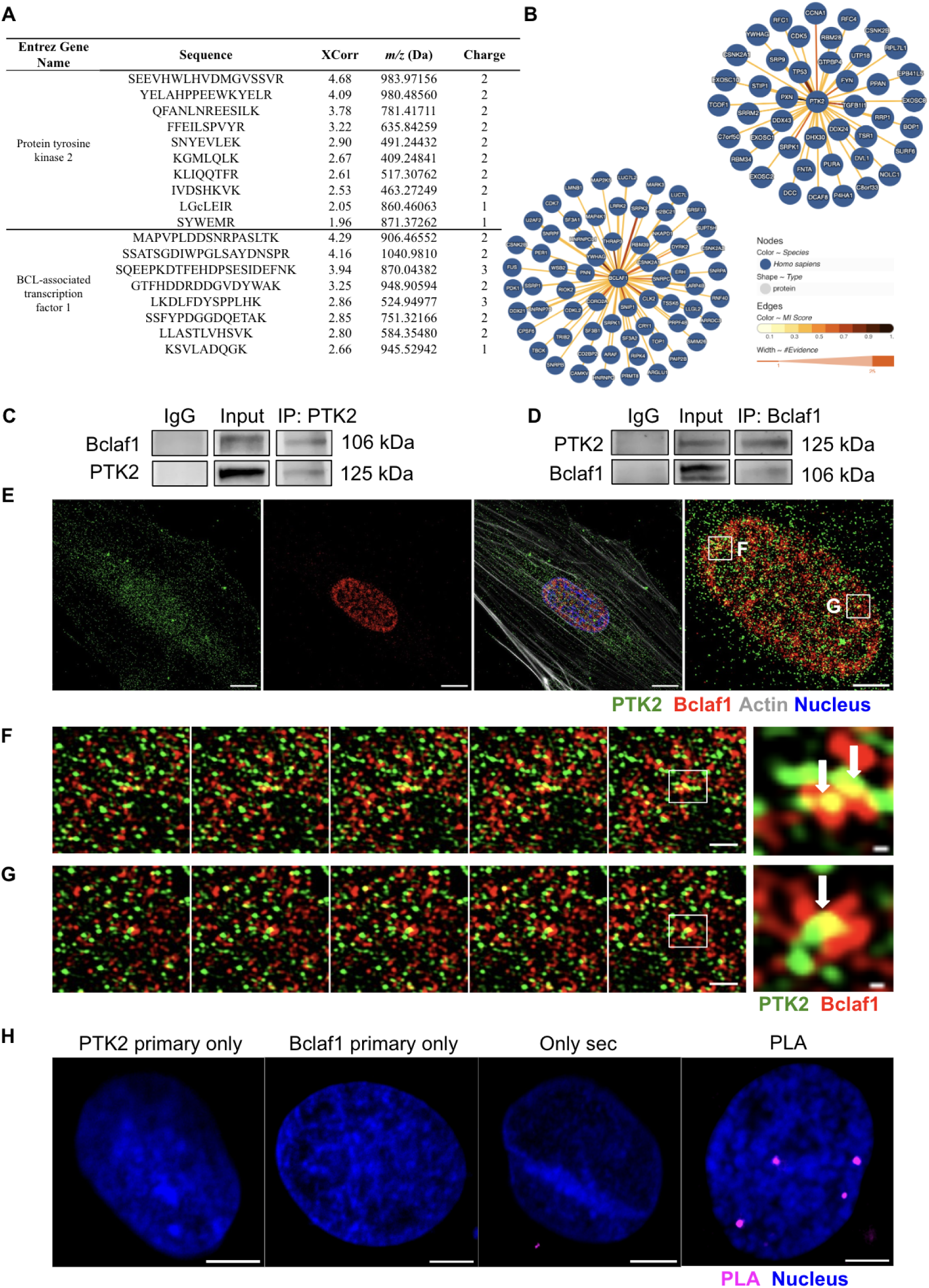
Bclaf1 is a binding partner of PTK2. **(A)** Partial results of mass spectrometry analysis of HEK-293T cells demonstrating the presence of Bclaf1 in Flag-FERM-PTK2 immunoprecipitated. Complete mass spectrometry data are provided in Supplementary Table 1. (**B)** Interaction networks of PTK2 and Bclaf1 on HEK293T obtained from IntAct (https://www.ebi.ac.uk/intact/home), including association and physical association. **(C-D)** Bclaf1 immunoblots of PTK2 immunoprecipitated from control H9c2 extracts confirming the PTK2-Bclaf1 interaction, and vice versa. PTK2 (C) and Bclaf1 (D) served as loading control. **(E)** Super-resolution images of H9c2 cells on basal conditions, labeled for PTK2, Bclaf1, actin, and nucleus. Scale bar = 10 µm. Nucleus zoomed image scale bar = 5 µm. **(F-G)** Z-stack slices of nuclear planes showing the association of PTK2 with Bclaf1 (arrows). Scale bar = 1 µm. Zoomed images scale bar = 0.1 µm. **(H)** Proximity Ligation Assay (PLA) demonstrating the interaction between PTK2 and Bclaf1 in untreated cells. Scale bar: 5 µm. Green: PTK2; Red: Bclaf1; Blue: nucleus; Gray: actin; Magenta: PLA.

Next, to explore the subcellular distribution of PTK2 and Bclaf1 in cardiomyocytes, super-resolution images were acquired by Structured Illumination Microscopy (SR-SIM). Figure 1E shows PTK2 distributed throughout the cytoplasm and nucleus and Bclaf1 mostly localized at the nucleus. On the right panel, overlapped channels show colocalization areas of PTK2 and Bclaf1. Two regions were highlighted, and Z-stack analysis showed evidence of PTK2-Bclaf1 interactions in the nuclear compartment of H9c2 cardiomyocytes (Fig. 1F-G). It was noticed that PTK2 and Bclaf1 displayed a diffuse distribution in the nucleus and colocalized in multiple points through the examined stacks. Proximity Ligation Assay (PLA) confirmed the PTK2-Bclaf1 direct interaction (Fig. 1H) establishing Bclaf1 as a new PTK2 interactor.

### 2.2 PTK2 and Bclaf1 are responsive to oxidative stress generated by doxorubicin, forming nuclear condensates

To investigate whether PTK2 and Bclaf1 are sensitive to oxidative stress (OS) generated by doxorubicin (dox), H9c2 cells were treated with this compound (12h; 1 µM) and analyzed by SR-SIM and western blotting. Dox treatment induced PTK2 nuclear accumulation and a rearrangement of Bclaf1 and PTK2 into prominent nuclear biomolecular condensates (Fig. 2A-B), which are non-membrane-bound biological structures with a physically defined boundary generated by surface tension^20^. Furthermore, we observed that dox treatment induces PTK2 autophosphorylation at tyrosine 397 residue, an essential event to switch PTK2 to an open and active conformation, culminating in PTK2 activation in the cell nuclei (Fig. 2C-D). Analysis of cytosolic and nuclear extracts confirmed the subcellular distribution of PTK2 and its activation in the nuclear compartment (Fig. 2E-F; Supplem. Fig. 7B-C). The autophosphorylation of tyrosine 397 induces PTK2 conformational changes which culminates in the exposition of protein-protein interactions sites and in the kinase domain activation^21^. These data indicate that PTK2 and Bclaf1 are sensitive to dox treatment and may regulate nuclear events to modulate cell behavior and fate during OS stimuli.

**Figure 2.**
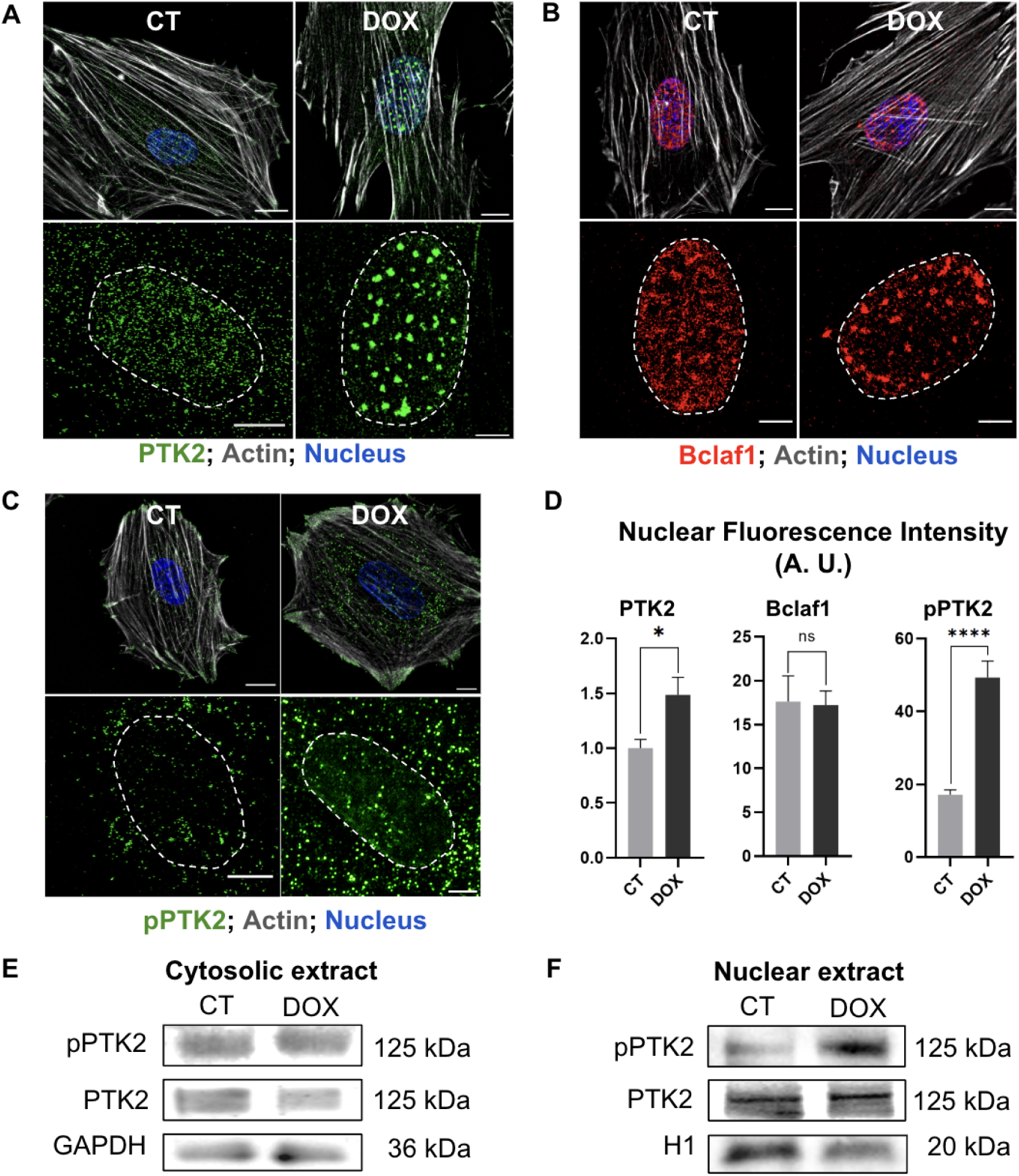
PTK2 and Bclaf1 are responsive to stress induced by dox. **(A-B)** Immunofluorescence images of H9c2 cardiomyocytes showing PTK2 and Bclaf1 reorganization in dot-like structures after dox treatment, respectively. **(C)** Immunofluorescence images of pY397-PTK2 (pPTK2) showing nuclear PTK2 activation after dox treatment. Scale bar = 5 µm. Scale bar of zoomed images = 3 µm. **(D)** Bar graphs show the nuclear fluorescence intensity of PTK2, Bclaf1, and pPTK2 on basal conditions and after dox treatment. Asterisks means statistical significance, while ns means P > 0.05; * means P ≤ 0.05; ** means P ≤ 0.01; *** means P ≤ 0.001 and **** means P ≤ 0.0001. **(E-F)** pY397-PTK2, PTK2, GAPDH, and Histone H1 (H1) specific immunoblots of cytoplasmic and nuclear extract of H9c2 control or treated with dox. Dox treatment: 1 µM; 12h. Green: PTK2 and pPTK2; Gray: Actin; Red: Bclaf1; Blue: nucleus.

We next examined whether the PTK2-Bclaf1 interaction persists during dox-induced stress. Co-IP assays revealed an association of Bclaf1 to the PTK2 immunoprecipitate, and vice versa (Fig. 3A-B; Supplem. Fig. 7A), confirming the PTK2-Bclaf1 interaction in H9c2 cardiomyocytes under dox-induced OS. Super-resolution images showed a high colocalization pattern of PTK2 and Bclaf1 nuclear condensates (Fig. 3C-D), indicating a very strong interaction. Even in non-dot-like regions PTK2 and Bclaf1 co-localize with a diffuse pattern (Fig. 3E). Furthermore, because our image resolution is ∼80-100 nm, it was possible to resolve the internal structure of these condensates (Fig. 3D-F), whose areas ranged from 200 to 700 nm². Line profile plot and 3D images showed Bclaf1 condensates may present a cavity containing PTK2, which points to an isolation of PTK2 inside the Bclaf1 condensate (Fig. 3C-F). Moreover, differentiated cardiomyocytes were able to form Bclaf1 condensates containing PTK2 when treated with dox (Supplem. Fig. 2A-C). Interestingly, PTK2-Bclaf1 biomolecular condensates exhibited different sizes and morphologies, from simple spherical shapes to complex structures with non-spherical topologies, which may reflect the presence of multiphase structures or different stages in their maturation^20,22,23^. Finally, proximity ligation assays (PLA) confirmed the PTK2-Bclaf1 direct interaction in dox-treated cardiomyocytes (Fig. 3G). Our results showed that PTK2-Bclaf1 nuclear condensates formed in response to OS in both precursor and differentiated cardiomyocytes.

**Figure 3.**
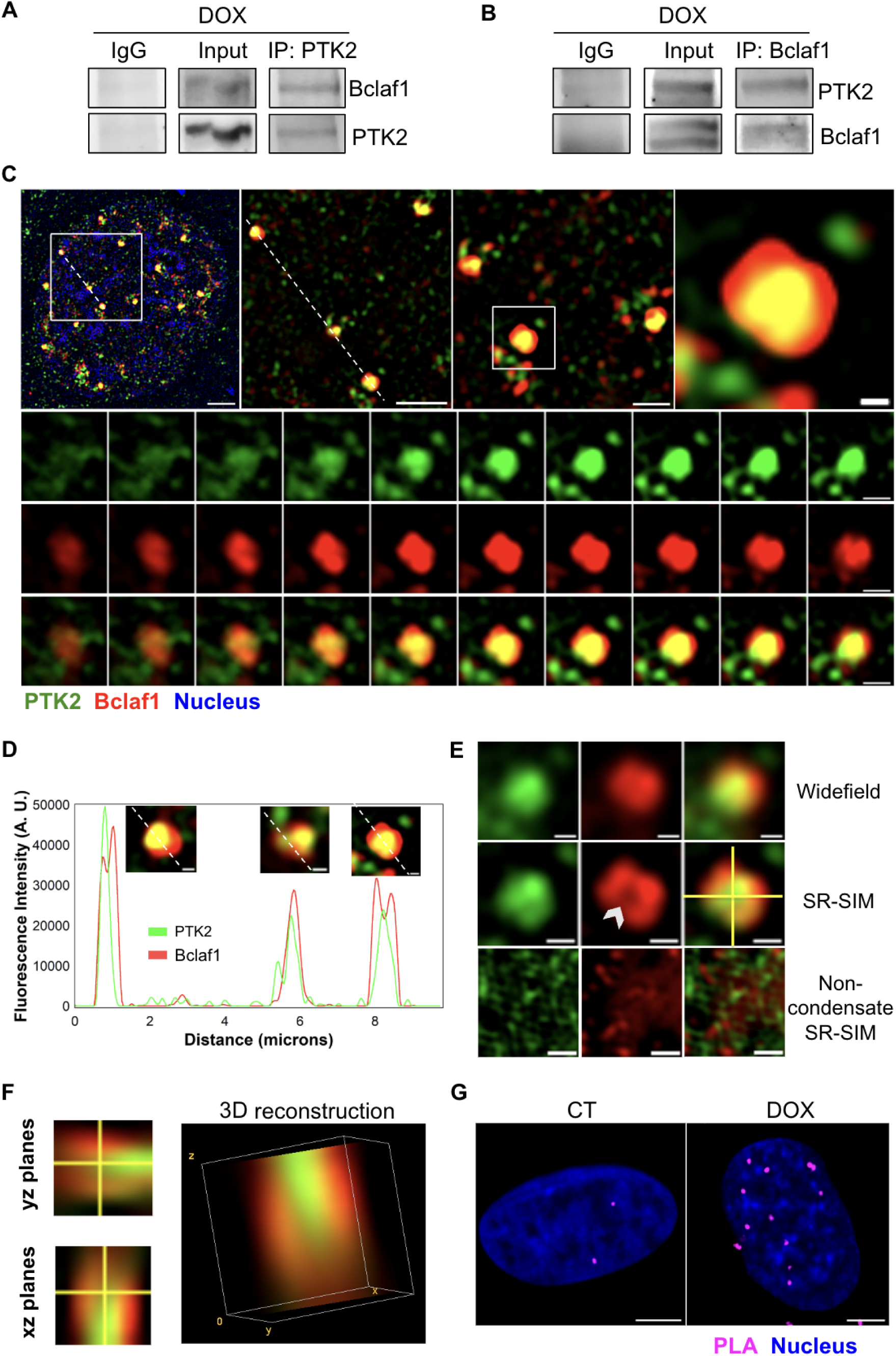
Oxidative stress induces PTK2 and Bclaf1 association into nuclear biomolecular condensates. **(A-B)** Bclaf1 immunoblots of PTK2 immunoprecipitate from dox-treated H9c2 cardiomyocytes extracts, and vice versa. PTK2 (A) and Bclaf1 (B) served as input loading control. **(C)** Immunofluorescence images of dox-treated cardiomyocytes showing the rearrangement of Bclaf1 into nuclear condensates containing PTK2. Below, Z-stack from nuclear planes, demonstrating the association of PTK2 and Bclaf1 in different planes. Scale bar of upper panels = 5 µm, 1 µm, 2 µm, and 0.5 µm, respectively. Scale bar of Z-stack images = 0.5 µm. **(D)** Fluorescence line profile plot of the horizontal line in (C) showing PTK2 inside Bclaf1 condensates. Scale bar = 0.5 µm. (**E)** Widefield and SR-SIM microscopy in a PTK2-Bclaf1 condensate, showing strong colocalization in contrast with a no-condensate region. Arrowhead shows a central cavity in the condensate. Scale bar = 0.5 µm. (**F)** Orthogonal Z projection showing yz planes, xz planes and 3D reconstruction of Bclaf1 biomolecular condensate containing PTK2 shown in “E”. Green: PTK2; Red: Bclaf1; Blue: nucleus. Dox treatment: 1 µM; 12h. **(G)** Proximity Ligation Assay showing increased interaction between PTK2 and Bclaf1 after dox treatment. Scale bar = 5 µm.

### 2.3 PTK2 interacts with Bclaf1 via liquid-liquid phase transition

To better characterize the PTK2-Bclaf1 interaction we employed structural model predictions. AlphaFold 3^24,25^ analysis revealed Bclaf1 as an intrinsically disordered protein (IDP) presenting an inner core of 4 alpha helices with flexible long loops surrounding it (Fig. 4A). The Predict Alignment Error (PAE) reflects this spatial conformation by having low value errors only on its diagonal in the PAE matrix. The unfolded pattern predicted for Bclaf1 indicates its ability to easily accommodate the docking of partner proteins, which corroborate its stabilization function described previously for HIF1-α and PD-L1^16,17,26^. Otherwise, PTK2 presents a much more stable predicted structure (Fig. 4B). The three main domains, which had their individual structure characterized, present low PAE scores meaning their positions are accurate. There is an intrinsically disordered region (IDR) in the N-terminal portion of the FERM domain and a longer IDR linking Kinase and FAT domains, compressed by Q686 to S934 residues, containing one alpha-helix (Q805-S832) positioned in the interface between the kinase and FAT domains. Pursuing Bclaf1 and PTK2 models, we next used the docking tool of AlphaFold3 to predict the PTK2-Bclaf interacting-models (Fig. 4C; Supplem. Fig. 3). From this prediction we could see an intimate association between Bclaf1 and PTK2, where Bclaf1 flexible loops can involve the entire structure of PTK2. The molecular mechanisms by which Bclaf1 interacts and stabilizes protein partners is unknown^16,27^, however, the results obtained from AlphaFold3 combined with its ability to form nuclear punctate structures raised the hypothesis that Bclaf1 undergo liquid-liquid phase separation (LLPS) to form biomolecular condensates, which increase the mobility of client proteins. During LLPS, biomolecules condense from the surrounding cellular environment into distinct membraneless compartments. This process concentrates proteins within the condensates, effectively increasing their binding affinity and minimizing competition with other interactors^28,29^.

**Figure 4.**
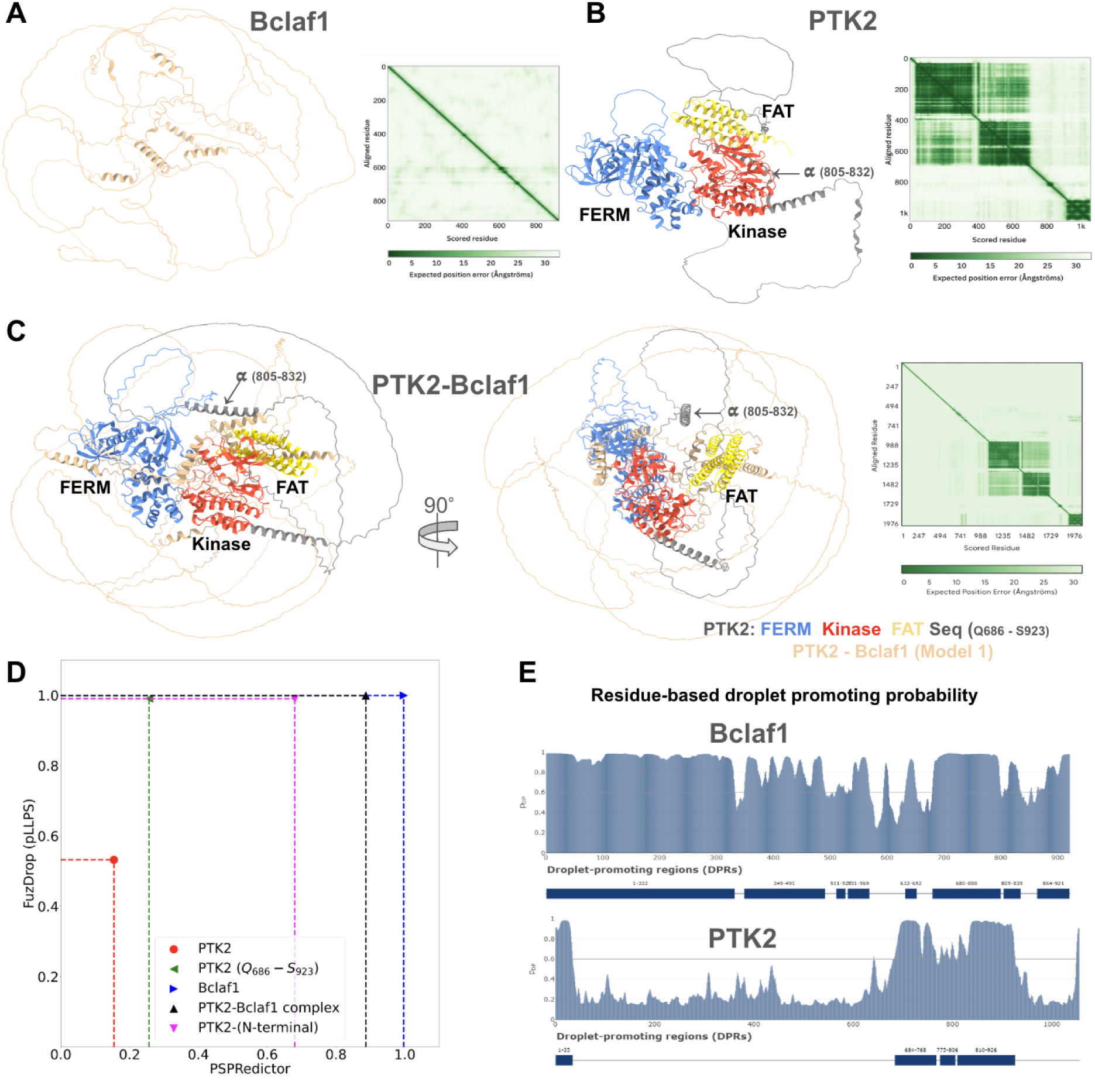
Computational predictions of PTK2 and Bclaf1 structure and interaction. **(A)** Structure of Bclaf1, an intrinsically disordered protein (IDP), with an inner core of 4 alpha helices and flexible long loops was obtained using AlphaFold3. PAE matrix indicates low error values only on the diagonal, reflecting the Bclaf1 unfolding nature. **(B)** PTK2 structure, with three main domains and a long intrinsically disordered region (IDR) linking the Kinase and FAT domains obtained by AlphaFold3 platform is shown. PAE matrix shows the predicted error in Angstroms. **(C)** Predicted PTK2-Bclaf1 model using AlphaFold3 docking tool and PAE matrix, indicating low error values for the PTK2 domains and a higher error value for the IDR. Interfaces of PTK2-Bclaf1 interaction with intermediate error values are shown. (**D)** LLPS propensity scores for Bclaf1, PTK2, IDR PTK2-N-terminal, Q686 - S923 IDR PTK2 sequence, and the PTK2-Bclaf1 complex obtained using FuzDrop and PSPredictor. **(E)** Residue-based droplet promoting probability of PTK2 and Bclaf1 obtained by FuzDrop.

To assess the propensity of Bclaf1 and PTK2, both as monomers and in complex, to undergo liquid-liquid phase separation, we calculated LLPS scores for Bclaf1, PTK2, and the PTK2-Bclaf1 complex using FuzDrop^27–29^ and PSPRedictor^30^ (Fig. 4D). As expected, the Bclaf1 scores were close to 1 (FuzDrop = 0.9995, PSPredictor = 0.9968) meaning it is prone to undergo LLPS. PTK2, as a monomer, presented lower scores (FuzDrop = 0.5332, PSPRedictor = 0.1543), indicating a low propensity for self-driven phase separation. Nevertheless, the intrinsically disordered N-terminal region and the linking between the Kinase and FAT domain has a high predicted propensity for phase separation according to the FuzDrop score (∼ 0.99), suggesting these regions might contribute to PTK2 LLPS when induced by an interacting partner. Finally, the scores for PTK2-Bclaf1 complex (PSPRedictor, FuzDrop) indicated a high probability to condensate via LLPS. The number of residues with a high probability (Predicted Drip Propensity > 0.5) of promoting liquid-liquid phase separation align with the data above (Fig. 4E). Although LLPS probability analysis indicates the presence of high-scoring residues in both PTK2 and Bclaf1, Bclaf1 exhibits a significantly higher number of these residues, suggesting it may function as the primary LLPS inducer for PTK2.

To validate the LLPS prediction data we performed Fluorescence Recovery after Photobleaching (FRAP) (Fig. 5A). When a protein is transitioned to LLPS its diffusion coefficient increases, reflecting a higher mobility in this phase^31,32^. In Figure 5B we plotted the average FRAP curves showing an increased diffusion of EGFP-Bclaf1 and EGFP-PTK2 in the condensate in contrast with the low mobility of these proteins outside the condensates, confirming that PTK2-Bclaf1 complex condensate via LLPS. We also performed Fluorescence-Lifetime Imaging Microscopy (FLIM) to further characterize the condensates^33^. The presence of distinct protein phases resulted in observable fluorescence lifetime variations. In our analysis, Bclaf1 exhibited an increased lifetime within condensates (1.095 ± 0.022 ns AVG ± SEM) when compared to nuclear regions (0.935 ± 0.062 ns). Similarly, PTK2 showed an increased lifetime in condensates (1.028 ± 0.026 ns) relative to non-condensate regions (0.854 ± 0.020 ns) (Fig. 5C-D). This data showed that Bclaf1 and PTK2 present higher mobility inside the biomolecular condensates and a different fluorescence lifetime as well, indicating a different phase for these proteins inside the condensates.

**Figure 5.**
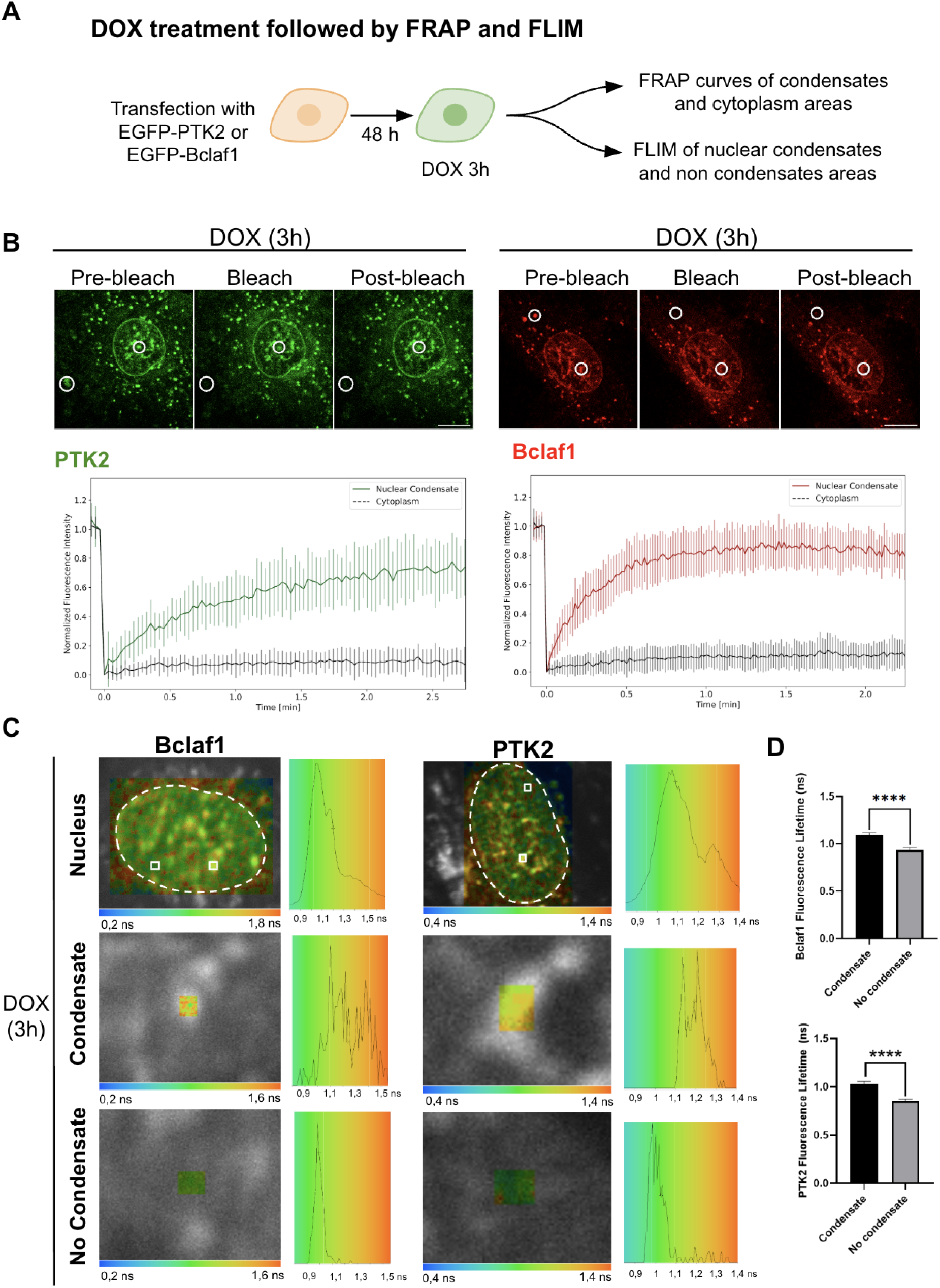
Bclaf1 and PTK2 undergo liquid-liquid phase separation to form submicron membraneless organelles. **(A)** Experimental design of FRAP and FLIM experiments. **(B)** FRAP images of condensate and cytoplasm regions of H9c2 cells transfected with EGFP-PTK2 and EGFP-Bclaf1. Scale bar = 10 μm. Below, FRAP curves are shown for PTK2 condensates (n=9) and cytoplasm regions (n=16) on the left, and for Bclaf1 condensates (n=11) and cytoplasm regions (n=11) on the right. **(C)** Lifetime images of H9c2 cells expressing EGFP-PTK2 or GFP-Bclaf1 fluorescent proteins. **(D)** Bar graph of EGFP-Bclaf1 (n = 4 cells) and EGFP-PTK2 (n = 6 cells) fluorescence lifetime in condensates and no condensate areas. Asterisks means statistical significance, while ns means P > 0.05; * means P ≤ 0.05; ** means P ≤ 0.01; *** means P ≤ 0.001 and **** means P ≤ 0.0001.

### 2.4 Hsp70 activity modulate the LLPS-mediated formation of PTK2-Bclaf1 condensates

Despite the recent studies on LLPS physical properties, the molecular chaperone role on the phase transition remains unclear. To verify whether Hsp70 activity modulates the dynamics of the PTK2-Bclaf1 biomolecular condensates, we performed SR-SIM and FRAP combined with chemical inhibition of Hsp70 (VER-155008; 50 µM) (Fig. 6A). Firstly, cells were incubated with dox for 1, 3, 6, and 12 h (1 µM) and analyzed by triple-labeling SR-SIM. As depicted on Figure 6B the clusterization of Hsp70 with PTK2 starts as early as 1 h of dox treatment. After 3 h, PTK2-Bclaf1 condensates were clearly visualized along with Hsp70 presence, following this same pattern over all the subsequent times, indicating an active participation of Hsp70 on their dynamic. Secondly, to confirm the Hsp70 contribution to this process, cells were examined by FRAP after being transfected with EGFP-Bclaf1 or with EGFP-PTK2 and co-treated with dox (3 h) and VER-155008 (1 h). Hsp70 inhibition reduced the diffusion of Bclaf1 and PTK2, as demonstrated by the lower recovery time after photobleaching (Fig. 6C). Together, our data indicated a possible role of Hsp70 in regulating the formation of biomolecular condensates, providing a deeper understanding of the dynamic behavior of these nuclear membraneless organelles during OS.

**Figure 6.**
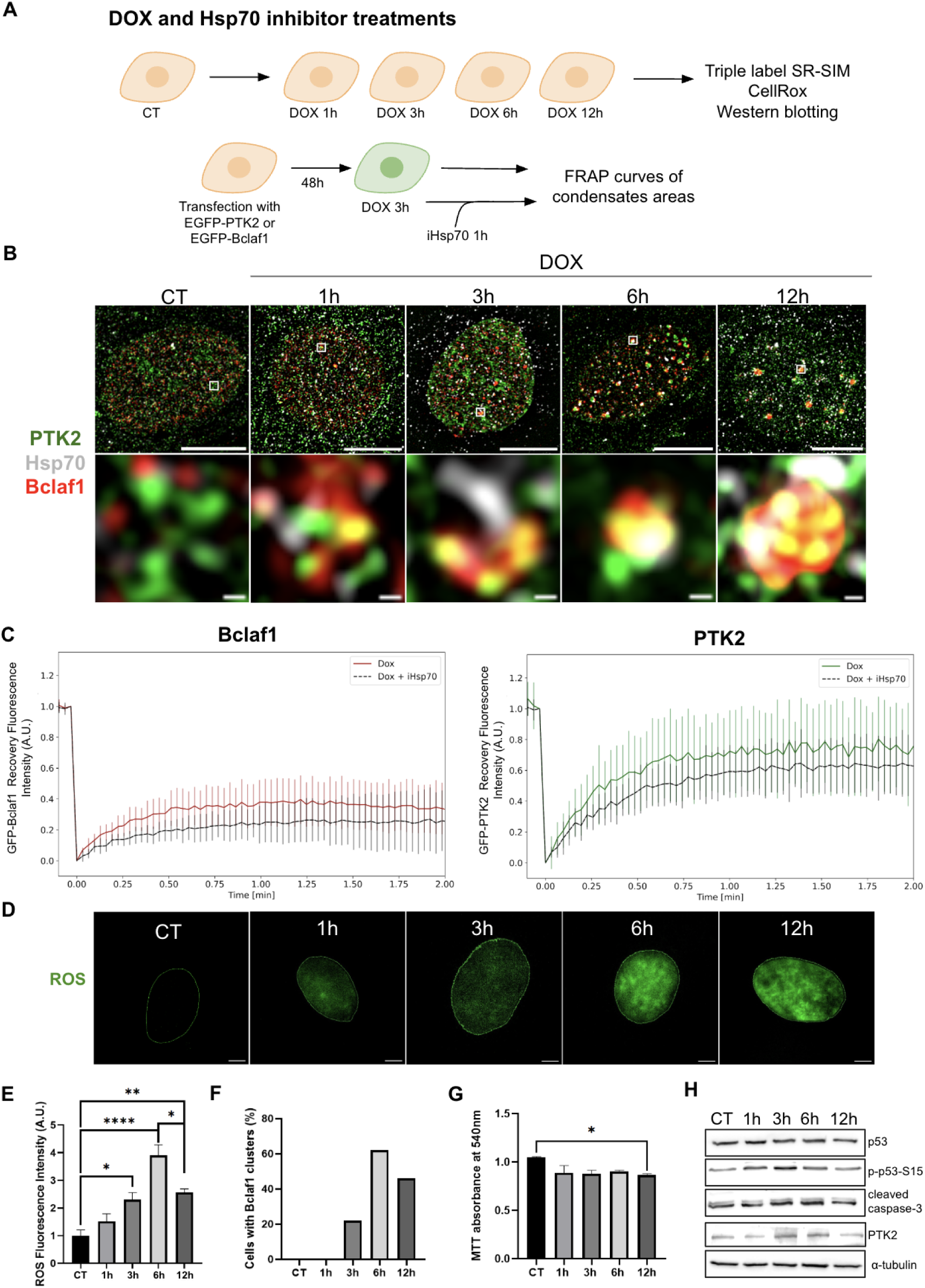
Hsp70 and oxidative stress modulates the condensate formation. **(A)** Experimental design of time-course and transfection experiments. **(B)** Super-resolution images of H9c2 cardiomyocytes showing the colocalization of Hsp70 (gray) with PTK2 (green) and Bclaf1 (red) on the condensate region. Scale bar = 10 µm. Scale bar of zoomed images = 0.5 µm. **(C)** FRAP curve showing the fluorescence recovery of EGFP-Bclaf1 (n = 9 regions for dox group, n = 8 for dox + iHsp70 group) and EGFP-PTK2 (n = 13 regions for dox and dox + iHsp70 group) on condensate areas after dox treatment followed Hsp70 inhibition (iHsp70). **(D)** Time-course experiments using CellROX green to detect and measure the production of ROS during dox treatment. Scale bar = 5 µm. **(E)** Bar graph shows CellROX fluorescence intensity during dox treatment (n = 5 cells for CT, n = 5 cells for 1h, n = 6 cells for 3h, n = 5 cells for 6h, and n = 5 cells for 12h). **(F)** Bar graph shows the percentage of cells with Bclaf1 clusters from control (CT) to 12h of dox treatment, as indicated (n = 80 cells for CT, n=59 cells for 1h, n = 94 cells for 3h, n = 76 cells for 6h, and n = 91 cells for 12h). **(G)** Bar graph shows the MTT absorbance at 540 nm. N = 7 replicates per group. **(H)** Western Blot images of p53, p53 phosphorylated at Ser 15 (p-p53-S15), and cleaved caspase-3. *α*-Tubulin served as loading control. Bar graphs are shown in Supplementary Figure 4.

### 2.5 The PTK2-Bclaf1 condensates formation correlated with the maintenance of basal p53 levels, contributing to cell survival

Next, we sought to analyze the relationship among PTK2-Bclaf1 biomolecular condensate formation, the generation of reactive oxygen species (ROS) and the activation of pro-apoptotic pathways. CellROX assays were performed on control and dox-treated cardiomyocytes. Cells were incubated for 1, 3, 6, and 12 h with dox and examined by fluorescence microscopy after the incubation with CellROX reagent (Fig. 6D). Image analysis showed a correlation of OS levels and the dynamic of biomolecular condensate formation. The level of ROS increased between 3 and 6 h of dox treatment, and started to reduce at 12 h (Fig. 6E). Interestingly, the PTK2-Bclaf1 condensate numbers follow the same pattern of increase as ROS levels, with a peak at 6 h and and a decrease at 12 h of dox treatment (Fig. 6F). MTT assays demonstrated a preservation of cell viability until 6 h of dox treatment, with a reduction after 12 h by ∼ 20 % in relation to control cells (Fig. 6G).

To investigate whether basal PTK2 levels were sufficient to maintain low p53 and suppress apoptosis pathways, we performed western blot assays for p53, phospho-p53, and cleaved caspase-3. These blots showed no significant alterations, with only a trend towards increased p53 observed after 12 h of dox treatment (Fig. 6H; Supplem. Fig. 4A; Supplem. Fig. 7D). This maintenance of p53 levels suggests that stabilized PTK2 was sufficient to limit p53 elevation, thereby promoting cell survival. Taken together, these data indicate that Bclaf1 and PTK2 undergoing LLPS is a cellular response to dox-induced oxidative stress, serving as a mechanism to maintain cell survival.

### 2.6 Bclaf1 biomolecular condensates protects PTK2 from the ubiquitin-proteasome system during dox-induced oxidative stress

Oxidative stress occurs when there is an imbalance between the production of reactive oxygen species and the intracellular antioxidant defenses, which can lead to potential damage, destabilizing proteins and other cell components^34,35^. As biomolecular condensates function in stress conditions by sequestering and stabilizing proteins^28,36,37^, we reasoned that Bclaf1 condensates were likely to be molecularly relevant to stabilize and protect PTK2 from aggregation and ubiquitination in the context of OS. To explore this hypothesis, we examined whether PTK2 localized into the Bclaf1 biomolecular condensate is less susceptible to ubiquitination. Triple-staining SR-SIM and image analysis showed that PTK2 diffusely disposed throughout the nucleus colocalizes with ubiquitin dot-staining, indicating the action of the ubiquitin-proteasome system (UPS) in sorting and degradation of PTK2 during dox-induced OS (Fig. 7A; panel 1). However, PTK2 localized within the Bclaf1 condensates showed very low ubiquitination (Fig. 7A; panel 2), indicating a role of the condensates to protect PTK2 from the UPS during OS. Next, we calculated the correlation of the integrated fluorescence of each channel. After automatic delimitation and identification of the Bclaf1 condensates area on Fiji software, a Python code calculated the Pearson correlation coefficient for the integrated fluorescence between PTK2, Bclaf1, and Ubiquitin channels. Higher values of this coefficient indicate the presence of both proteins at a given region, while negative values indicate they are anticorrelated and zero indicates no correlation. Figure 7B shows that PTK2 and Bclaf1 have a higher correlation coefficient (0.89 ± 0.02; AVG ± SEM) than PTK2 and Ubiquitin (0.68 ± 0.05). This suggests that PTK2 becomes less susceptible to ubiquitination when sequestered in the biomolecular condensate.

**Figure 7.**
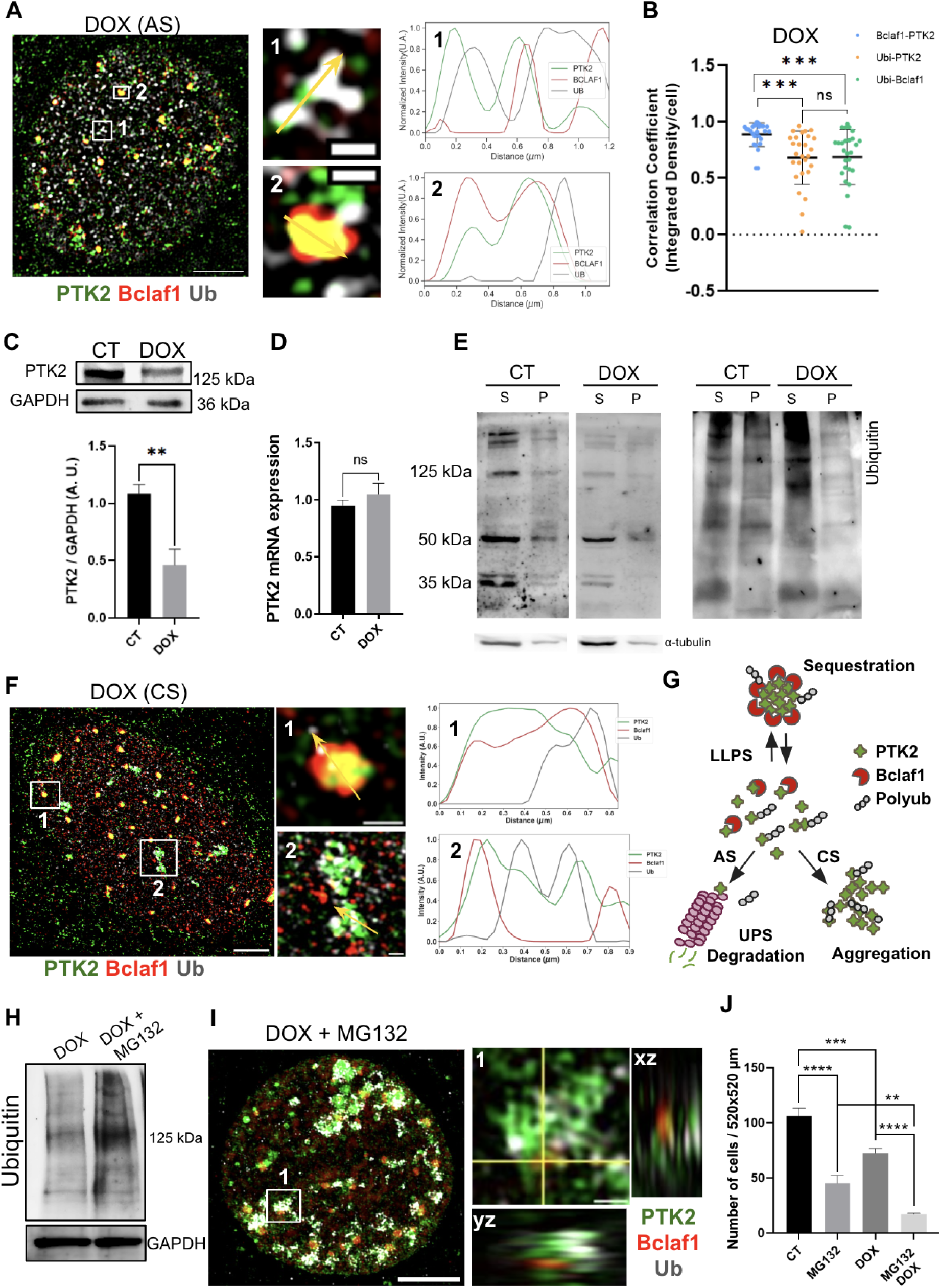
Bclaf1 biomolecular condensates protect PTK2 from UPS-mediated ubiquitination. **(A)** Triple-label SR-SIM image showing PTK2 and Bclaf1 concentrated on biomolecular condensates on the nucleus of H9c2 cardiomyocytes treated with dox. Scale bar = 10 µm. **1.** Fluorescence line profile plot showing PTK2 colocalizing with ubiquitin outside the Bclaf1 condensate. Scale bar = 0.5 µm. **2.** Fluorescence line profile plot showing low ubiquitin level inside the Bclaf1 condensate, where PTK2 is enriched. Scale bar = 0.5 µm. **(B)** Pearson correlation coefficient between Bclaf1 and PTK2, ubiquitin and PTK2, and ubiquitin and Bclaf1. **(C)** PTK2 immunoblots of extracts from control (CT) and dox-treated (DOX) H9c2 cells and bar graph showing a decrease in PTK2 levels after dox treatment. N = 3 biological replicates per group. **(D)** RT-qPCR assay shows no difference in PTK2 mRNA levels in cardiomyocytes on basal conditions or after dox treatment. GAPDH was used as endogenous control. N = 4 biological replicates per group. **(E)** Western-blot of PTK2 and ubiquitin comparing soluble (S) and insoluble (P) fractions of H9c2 cardiomyocytes. **(F)** Super-resolution images of PTK2, Bclaf1, and ubiquitin staining after 36 hours of recovery from dox treatment. Scale bar = 5 µm. 1. Fluorescence line profile plot showing low PTK2 ubiquitination inside Bclaf1 condensate. Scale bar = 0.5 µm. 2. Fluorescence line profile plot showing PTK2 ubiquitinated and agglomerated outside Bclaf1 condensate. **(G)** Graphical representation of PTK2-Bclaf1 undergoing LLPS to avoid PTK2 ubiquitination and aggregation in cells under oxidative stress. Bclaf1 sequesters PTK2 into biomolecular condensates in cells submitted to stress. Under acute stress (AS), PTK2 outside the condensate is primary ubiquitinated and targeted for proteasomal degradation, while under chronic stress (CS), ubiquitinated PTK2 accumulates in nuclear aggregates. **(H)** Western-blot showing ubiquitin accumulation on cardiomyocytes treated with dox and co-treated with dox and MG132. **(I)** SR-SIM image showing the accumulation of ubiquitinated PTK2 on the nucleus of cardiomyocytes co-treated with dox and MG132. Scale bar of nuclear image = 5 µm. **(1)** Orthogonal Z projection showing yz planes and xz planes of a PTK2 agglomerate, which colocalize with ubiquitin. Scale bar = 0.5 µm. **(J)** Cell number count after dox combined with MG132 treatment, as indicated. N = 15 images of each group. Green: PTK2; Red: Bclaf1; Gray: Ubiquitin. Asterisks demonstrates statistical significance, while ns means P > 0.05; * means P ≤ 0.05; ** means P ≤ 0.01; *** means P ≤ 0.001 and **** means P ≤ 0.0001.

As diffuse PTK2 may present a higher chance of being ubiquitinated, we used western blotting and RT-qPCR assays to investigate whether PTK2 levels were reduced after dox-induced OS. Accordingly, our data confirmed a reduction in PTK2 protein levels in 12h dox-treated cells (Fig. 7C; Supplem. Fig. 7E), while RT-qPCR analysis revealed no significant change in PTK2 mRNA levels (Fig. 7D), pointing to an increase in PTK2 degradation. To analyze PTK2 quality control during dox-induced OS, soluble (S) and insoluble (P) extracts were prepared. Western blot analysis of S and P samples from control and dox-treated cells demonstrated the presence of the 125 kDa band, corresponding to PTK2 Full Length (PTK2-FL) and the characteristic proteolytic fragments derived from calpain or caspase 3 cleavage^4,36^ (Fig. 7E). After dox treatment, the abundance of PTK2-FL in the soluble fraction decreased, while the proportion of the 50 kDa fragment increased, suggesting increased fragmentation of PTK2 during dox exposure. Moreover, the presence of PTK2 and its fragments on soluble fractions indicates that they are readily accessible for ubiquitination and UPS-mediated degradation. Corroborating this hypothesis, dox-treated cells exhibited increased ubiquitination in the soluble fraction (Fig. 7E), supporting that dox-mediated PTK2 downregulation is a consequence of ubiquitination and subsequent proteolytic degradation. Taken together, these findings indicate that dox-induced OS leads to PTK2 ubiquitination and degradation, while Bclaf1 condensates counteract this process by protecting nuclear PTK2 from oxidative damage and subsequent UPS activity.

To get more information about PTK2 stability and ubiquitination after dox treatment, H9c2 cells were treated with dox for 12h and incubated for 36h in a dox-free medium resulting in an extended exposition to OS (chronic OS) (Supplem. Fig. 5). SR-SIM analysis demonstrated the formation of PTK2-Bclaf1 condensates and the accumulation of ubiquitinated-PTK2 aggregates within the nuclear compartment under chronic OS, indicating the sustained effects of oxidative damage culminating in PTK2 destabilization and ubiquitination (Fig. 7F). Our results suggest a dynamic mobility of PTK2 to the Bclaf1 biomolecular condensates, where this kinase is protected from the UPS action. However, PTK2 diffused through the nucleus can be destabilized by OS resulting in its ubiquitination and degradation during acute OS or in its accumulation in amorphous nuclear aggregates in cells submitted to chronic OS, indicating a UPS overload (Fig. 7G). Next, to better understand the control of PTK2 proteostasis, H9c2 cardiomyocytes were treated with the proteasome inhibitor MG132 to induce a disruption of UPS-dependent proteostasis during dox-induced OS. Western blot analysis shows an increase in the overall ubiquitination after MG132 treatment (Fig. 7H). As expected, our super-resolution images showed an extensive accumulation of ubiquitinated PTK2 in the nuclei (Fig. 7I). Additionally, Bclaf1-biomolecular condensates were abundant after the inhibition of the proteasome, indicating Bclaf1 LLPS increased after the co-treatment with dox and MG132. However, the physical characteristics of these condensates were not assessed. In Figure 7I, a zoomed nuclear region is demonstrating ubiquitin presence within PTK2 aggregates and no colocalization with Bclaf1 biomolecular condensates (panel 1). This confirms that UPS degradation is involved in the clearance of PTK2 that is not sequestered and stabilized into the condensates. Furthermore, a prominent cell death was observed for cardiomyocytes co-treated with dox and MG132 (Fig. 7J), reflecting the severe effect of impaired proteostasis combined with oxidative damage.

Our SR-SIM data revealed that disrupting proteostasis through proteasome inhibition results in significant PTK2 ubiquitination and accumulation in nuclear amorphous aggregates, further supporting the role of the UPS in regulating PTK2 degradation. On the other hand, previous studies had not yet demonstrated an ubiquitination site on PTK2 structure. To verify whether PTK2 is directly ubiquitinated, we performed Co-IP coupled to mass spectrometry using H9c2 cardiomyocyte extracts. This approach enabled us to identify an ubiquitination site of PTK2 located on lysine 926 residue (Fig. 8A). In agreement with our findings, the Phosphosite Platform predicts that the peptide containing the K926 residue is a potential target for ubiquitination^38^ (Supplem. Fig. 6). This residue is located at the beginning of the FAT domain, as shown in the linear representations of PTK2-Rat and PTK2-Human (Fig. 8B). Notably, the PTK2 amino acid sequence is highly conserved between rats and humans (Supplem. Fig. 6B), with K926 in rats being equivalent to the K923 residue in humans. The K926 site is accessible on the FAT-PTK2 structure (Fig. 8C), making it a potential target for ubiquitin ligase complexes. Computational modeling of the PTK2-Bclaf1 interaction predicts that Bclaf1 may prevent K926 ubiquitination through steric occlusion of the site during interaction with PTK2 (Fig. 8D). Moreover, our experimental data indicates that within biomolecular condensates, Bclaf1 molecules assemble into molecular layers effectively shielding PTK2. Together, our findings confirm PTK2 as a target of the UPS, underscoring the role of the UPS in regulating PTK2 proteostasis during dox-induced OS. Furthermore, the sequestration of PTK2 by Bclaf1 biomolecular condensates represents a higher-level regulatory mechanism that modulates the activity of the UPS on the degradation of PTK2, a pro-survival kinase, during stress conditions.

**Figure 8.**
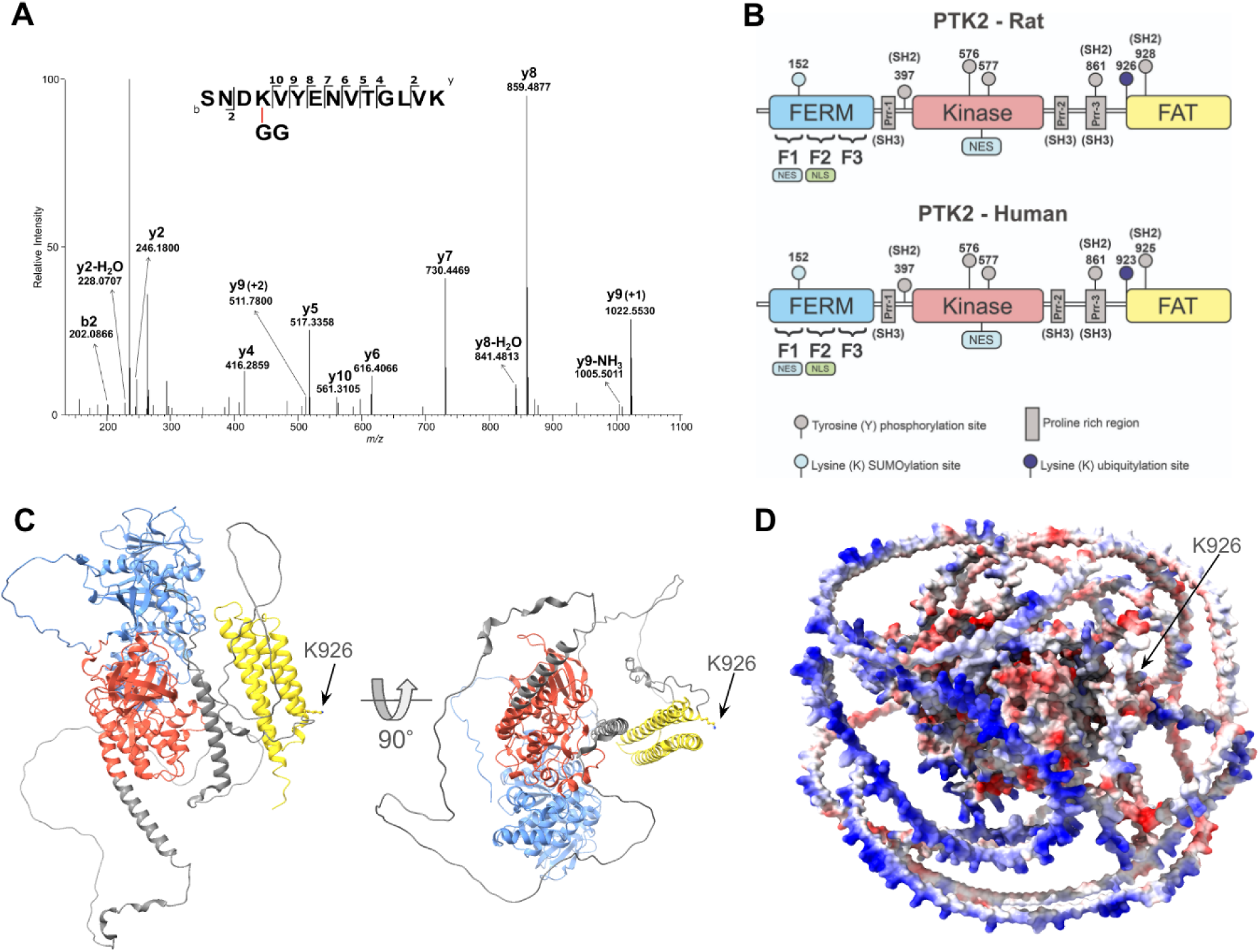
PTK2 is ubiquitinated on lysine 926 residue. **(A)** MS/MS spectrum showing the PTK2 identified peptide (m/z 561.3040) with the chemical modification (K-GG) characteristic of ubiquitination. **(B)** Linear representation of rat and human PTK2 demonstrating their domains and the PTK2 ubiquitination site on rat, lysine 926, and in human, lysine 923. **(C)** Cartoon representation of PTK2 showing the K926 ubiquitination site on the FAT domain. **(D)** Electrostatic surface representation of PTK2-Bclaf1 interaction complex, showing K926 site being sterically occluded by Bclaf1. The FERM domain is represented in blue, the Kinase in red, and the FAT domain in yellow.

### 2.7 Bclaf1 biomolecular condensates regulate PTK2 stability to enhance cell survival

To confirm the role of Bclaf1 on PTK2 stabilization on the nucleus of cardiomyocytes during dox-induced oxidative damage, shRNA-gene knockdown and super-resolution microscopy experiments were performed. Gene knockdown (KD-Bclaf1) was confirmed by western blotting experiments (Fig. 9A; Supplem. Fig. 7F). Surprisingly, KD-Bclaf1 cells treated with dox presented a significant increase in overall protein ubiquitination, accessed by western blotting, in relation to dox-treated control cells (CT) (Fig. 9B). The increased level of protein ubiquitination after Bclaf1 depletion was confirmed by SR-SIM analysis (Fig. 9C). As expected, in KD-Bclaf1 dox-treated cells, the Bclaf1 transition to liquid phase was disrupted (Fig. 9D). Moreover, nuclear PTK2 becomes organized into amorphous agglomerates colocalizing with ubiquitin staining, suggesting protein destabilization and ubiquitination due to the absence of Bclaf1 biomolecular condensates during dox-induced OS (Fig. 9E). To confirm whether PTK2 would be destabilized and tagged to UPS degradation in dox-treated KD-Bclaf1 cells, we analyzed PTK2 expression by western blot and RT-qPCR. Our data showed a reduction of PTK2 protein levels in KD-Bclaf1 dox-treated cells (Fig 9F; Supplem. Fig. 7G). In contrast, PTK2 mRNA levels were increased (Fig. 9F), suggesting an increase in PTK2 degradation and an activation of a transcriptional response possibly trying to keep the basal levels of PTK2. The reduction of PTK2 amounts can be attributed to the absence of Bclaf1 condensates protecting this kinase from the UPS. These data sustained a molecular mechanism in which Bclaf1 biomolecular condensates sequesters and protects PTK2 from oxidative damage, preventing its destabilization and UPS degradation during dox treatment.

**Figure 9.**
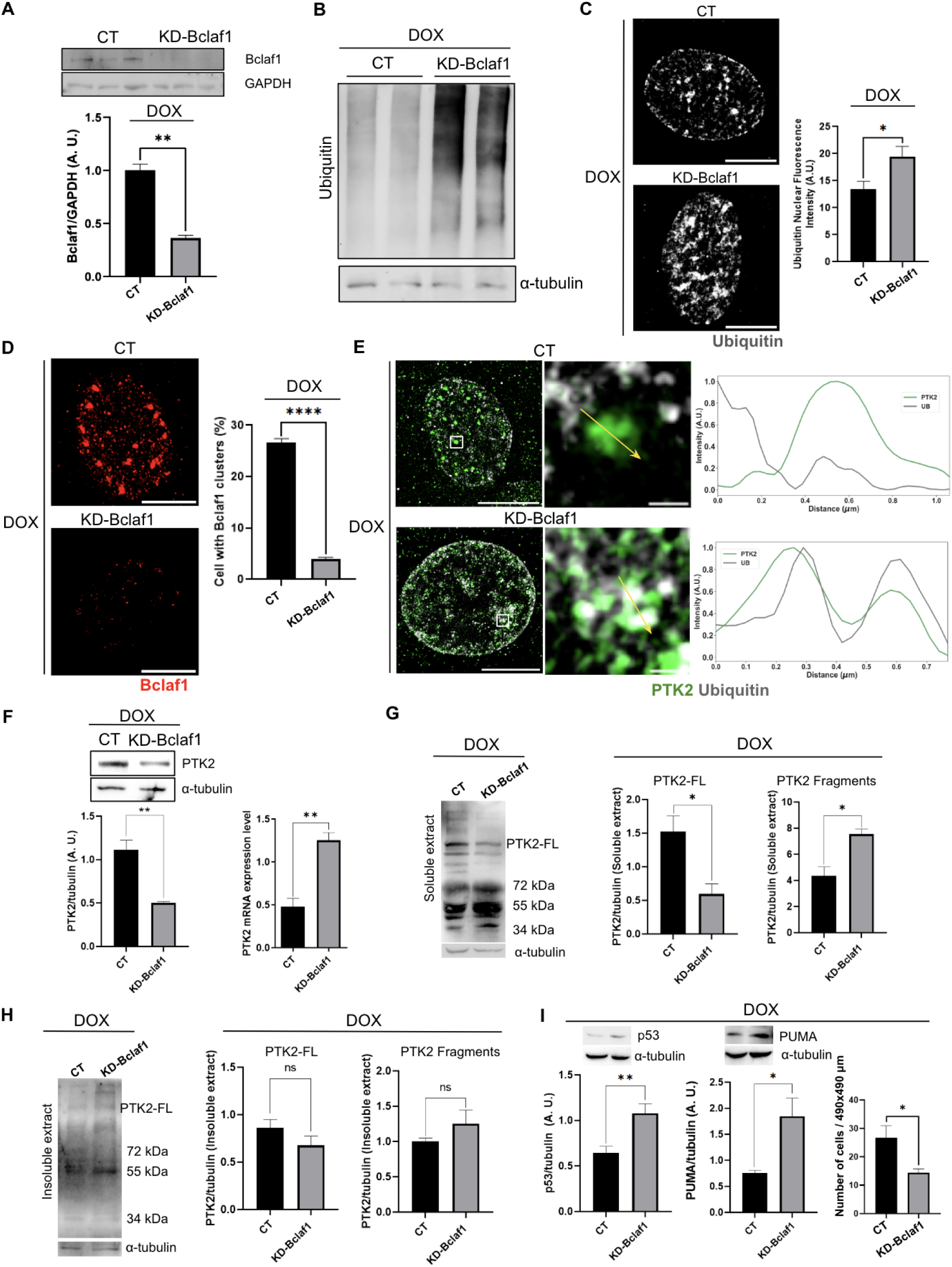
Bclaf1 knockdown results in PTK2 degradation and apoptosis activation. **(A)** Western blot image and bar graph confirming the knockdown of Bclaf1 (KD-Bclaf1) in relation to control (CT) wild type H9c2 cardiomyocytes. N = 3 biological replicates per group. **(B)** Western blotting images show the increased ubiquitination after Bclaf1 knockdown in relation to control cardiomyocytes, both treated with dox. **(C)** SR-SIM image and bar graph show the ubiquitin accumulation on the nucleus of KD-Bclaf1 cardiomyocytes treated with dox. Scale bar = 5 µm. N=24 cells per group. **D)** SR-SIM image shows the effect of Bclaf1 knockdown on disrupting condensates assembly. Scale bar = 5 µm. N=321 cells of Empty group and N=173 cells of group KD-Bclaf1. **(E)** SR-SIM image and line profile plot showing Bclaf1 knockdown increases PTK2 agglomeration and ubiquitination on the nucleus. Scale bar = 5 µm. Scale Bar of zoomed image = 0.5 µm. 1. Fluorescence line profile plot showing low PTK2 ubiquitination in PTK2 condensate. 2. Fluorescence line profile plot showing PTK2 ubiquitinated outside the condensate. **(F)** Western Blot and bar graphs show PTK2 levels reduction on KD-Bclaf1 cells treated with dox, while PTK2 mRNA expression levels increased. N = 4 biological replicates per group. **(G)** Western blot of soluble extracts from control and KD-Bclaf1 H9c2 cells treated with dox. Bar graphs show a reduction of PTK2-Full length (PTK2-FL) band and an increase in PTK2 cleaved bands (55 KDa and 34 kDa). **(H)** Western blot and bar graph analysis of insoluble extract from control and KD-Bclaf1 H9c2 cells treated with dox. The analysis showed no difference between CT and KD-Bclaf1 cells on the amount of PTK2-FL and PTK2 fragments present on the insoluble fraction. **(I)** Western blot and bar graphs show increased p53 and PUMA levels on KD-Bclaf1 H9c2 cardiomyocytes treated with dox. N = 3 biological replicates per group. On the right, the bar graph shows a reduced number of cells on the KD-Bclaf1 cells in relation to control cells, after dox treatment. N=321 cells of Empty group and N=173 cells of group KD-Bclaf1.

Supporting the role of Bclaf1 on PTK2 stabilization, Bclaf1 loss-of-function reduced soluble PTK2-FL and increased its fragmentation after dox treatment (Fig. 9G). Analysis of the insoluble fractions demonstrated no alterations on PTK2 bands pattern after Bclaf1 knockdown (Fig. 9H). These results corroborated our previous data demonstrating PTK2 fragments are primarily soluble and accessible for ubiquitination and degradation by the UPS. Furthermore, our data confirm Bclaf1 as a stabilizing protein that works, via LLPS, protecting its client proteins from the ubiquitination and degradation on cells under stress.

Given the well-established pro-survival role of PTK2^6,12,13,39^, we examined whether the degradation of PTK2 following Bclaf1 knockdown would affect cell survival. Our data demonstrated an accumulation of p53 and PUMA, pro-apoptotic proteins, on dox-treated KD-Bclaf1 cardiomyocytes (Fig. 9I; Supplem. Fig. 7H), indicating the activation of cell death by apoptosis. Finally, to verify the percentage of cell death on dox-treated KD-Bclaf1 cells, panoramic images were obtained to count the number of adhered cells in each field. The dox treatment was done after cells reach 80% of confluence and the analysis confirms the reduction of approximately 40% on cell number after 12 h of dox treatment in KD-Bclaf1 cells in relation to CT cells. These data link the stabilization of PTK2 by Bclaf1 condensates with the cell’s ability to resist dox-induced OS. Taken together, data obtained from Bclaf1 knockdown and proteasome inhibition revealed a new molecular mechanism whereby Bclaf1 biomolecular condensates stabilize and protect PTK2 from ubiquitination and degradation by the ubiquitin-proteasome system during dox-induced OS in cardiomyocytes, promoting cell survival.

## 3. DISCUSSION

Using a multi-faceted approach, we identified nuclear Bclaf1 biomolecular condensates being crucial for PTK2 stabilization during dox-induced oxidative stress. These condensates are submicron-scale membraneless organelles formed by LLPS that sequester and protect PTK2 from ubiquitin-proteasome-mediated degradation. Disruption of Bclaf1 condensates led to increased PTK2 ubiquitination and degradation, ultimately activating cell death via apoptotic pathways. We also identified a specific lysine ubiquitination site on PTK2, providing the first experimental evidence of ubiquitin-proteasome system involvement in regulating PTK2 levels in cardiomyocytes. Our findings establish Bclaf1 biomolecular condensates as nuclear sites that counteracts the activity of the ubiquitin-proteasome system over PTK2 during dox-induced OS, highlighting a novel protein quality-control mechanism that operates stabilizing proteins to modulate cell survival.

An important aspect of our Co-IP coupled to MS/MS data was the use of a PTK2 Flag-FERM-domain as a bait to identify new PTK2 interactors prone to stabilize PTK2 during stress. Consequently, PTK2 FERM domain is enough to mediate its interaction with Bclaf1, an essential process to enhance cell survival during OS. The PTK2 FERM mediates protein-protein interactions that are necessary for PTK2 function in the nuclear subcompartment^1^. In the nucleus, PTK2-FERM domain interacts with and regulates the levels of transcription factors, including p53 and GATA4^6^. FERM forms a complex with p53 and Mdm2, an ubiquitin ligase family member, bringing these proteins into close proximity and facilitating p53 ubiquitination and degradation via UPS, ultimately promoting cell survival^12^. In this study, we describe a novel pro-survival mechanism whereby PTK2 is stabilized and protected from UPS-mediated degradation by Bclaf1 biomolecular condensates during OS. The maintenance of functional PTK2 may consequently support its p53-Mdm2 regulatory pro-survival mechanism. Although the search for new nuclear interacting partners for PTK2 FERM-domain has been increasing, FERM mediating Bclaf1 interaction was described for the first time and adds a new layer of information on pro-survival mechanisms activated during stress.

Corroborating the responsiveness of PTK2 to different kinds of stimuli^2,4,8,40^, we demonstrated that PTK2 accumulates in the nucleus of cardiomyocytes under OS. Moreover, our data demonstrated that dox-induced OS led to the phosphorylation of tyrosine 397 of PTK2, a necessary event for PTK2 activation (Fig. 2C, F). Inactive PTK2 is found in an auto-inhibited conformation, with FERM binding to the Kinase domain, physically inhibiting its activation^21^. PTK2 activation requires dimerization, concomitantly with the binding of the FERM domain to phosphatidylinositol-4,5-biphosphate, allowing PTK2 Y397 autophosphorylation and activation^41^. Then, in the open conformation, binding sites are exposed to modulate signaling pathways through protein-protein interactions^41–43^. The responsiveness of PTK2 to dox-induced oxidative and, consequently, genotoxic stress accumulating in the nucleus of cardiomyocytes may be linked with its role in regulating p53 levels^6,12^. Indeed, when PTK2 levels were reduced after Bclaf1 knockdown, we observed increased p53 levels (Fig. 9I), ultimately resulting in apoptosis. In line with our findings, previous studies in mice have demonstrated that PTK2 protects the heart during dox treatment by maintaining appropriate p21 levels to promote cell survival^13^.

Using super-resolution microscopy and FRAP approaches, we demonstrated that PTK2 localizes within Bclaf1 biomolecular condensates in dox-treated cardiomyocytes (Fig. 3), supporting previous findings on condensate formation in response to stress^28,29,44^. Biomolecular condensates are membraneless compartments in which protein can be sequestered, modified, and stabilized^45–47^. Previous studies demonstrated Bclaf1 interacting with and stabilizing HIF1-*α* and PD-L1 by modulating their ubiquitination levels during hypoxia and after irradiation, respectively, although the underlying mechanisms remain poorly understood^16,17,48^. Nevertheless, no studies have examined the potential role of Bclaf1 in stabilizing proteins through LLPS during stress. Here, we propose an overall mechanism by which Bclaf1 counteracts the ubiquitin-proteasome system, through liquid-liquid phase separation, stabilizing proteins involved in adaptive stress responses, modulating cell fate. Bclaf1 is an intrinsically disordered protein^49^, whereas PTK2 contains two intrinsically disordered regions (Fig. 4E). Our analysis suggests that Bclaf1 primarily undergoes LLPS forming nuclear biomolecular condensates. On the other hand, the interaction of Bclaf1 with the PTK2 IDRs may induce PTK2 phase transition and translocation into the membraneless organelles, where this kinase is stabilized and protected from the UPS. FRAP experiments revealed enhanced mobility for both proteins <u>within</u> the condensates (Fig. 5B). FLIM results demonstrated increased fluorescence lifetimes in both Bclaf1 and PTK2 condensates (Fig. 5C), likely due to the increased concentration of these proteins into these structures. These findings align with previous observations in other membraneless organelles, including speckles, stress granules, and the nucleolus^50^. Interestingly, previous studies demonstrated that PTK2 undergoes LLPS in focal adhesions in the presence of paxillin. The droplet formation was linked to PTK2 oligomerization, facilitated by the interaction between FERM-FERM and FERM-FAT domains, which promoted the formation of liquid condensates on focal adhesions^50,51^.

Studies have demonstrated that molecular chaperones may regulate the biomolecular condensates physiology^29^. Chaperones are essential proteins that actively remodel protein-protein interactions and facilitate protein folding^52,53^. While their role in maintaining protein quality and preventing aggregation is well-established, the emerging field of biomolecular condensates is revealing their action in regulating the material properties and functions. For instance, disaggregases chaperones, including Hsp70 and AAA+ATPases, can modify the porosity of condensates, facilitating a quicker exchange of materials with the surrounding environment^29,54^. Chaperones can also have a role in condensate assembly or disassembly^28^. In this context, using FRAP experiments combined with Hsp70 inhibition, we confirmed the impact of Hsp70 activity on the Bclaf1 mobility into early formed biomolecular condensates (Fig. 6B-C). Some studies exploring Hsp70 role on condensate dynamics suggest an action for this chaperone on the quality control and maintenance of the condensates^55^, while others suggest chaperones promote the disassembly of the condensates, removing proteins during stress recovery^37^. Our data indicates a role of Hsp70 on the dynamics of Bclaf1 condensates, possibly restructuring their components to ensure their continued activity during dox-induced OS.

Oxidative stress generated by dox is a primary cause of cardiotoxicity^56^. The overproduction of ROS induces oxidative damage to biological macromolecules, including proteins, lipids, and DNA, and can induce cardiomyocyte apoptosis^57,58^. Proteins are the primary targets of oxidative modifications, which induces conformational changes and the exposure of internal regions to the solvent, resulting in protein ubiquitination. Oxidatively modified proteins must be degraded by the proteasome system to maintain cellular proteostasis and, ultimately, cell survival^59,60^. Our data revealed a progressive increase in OS from 1 to 6 hours of dox treatment, reaching a peak at 6 hours. This coincided with the nuclear accumulation of Bclaf1 biomolecular condensates, suggesting Bclaf1 LLPS is a response to elevated ROS levels (Fig. 6C-G). Additionally, cell viability remained preserved up to 6 hours of treatment, indicating that LLPS plays a role in enhancing cell survival during OS. Next, in line with the effects of dox in the oxidation and ubiquitination of proteins, we demonstrated that PTK2 is susceptible to the effects of oxidative damage, becoming fragmented and targeted by the UPS after dox treatment (Fig. 7). Analysis of soluble and insoluble fractions confirmed the propensity of PTK2 to remain in the soluble fraction, prone to be degraded by proteasomes after 12 hours of dox treatment. Alternatively, ubiquitinated PTK2 can aggregate in the nucleus of cardiomyocytes exposed to 48 hours of chronic OS. Furthermore, PTK2 sequestered into the Bclaf1 condensates becomes less accessible to be ubiquitinated and, consequently, less degraded by proteasomes. These findings corroborate the previous function described for Bclaf1 in the stabilization of HIF-1a during hypoxia^17^.

The regulation of PTK2 homeostasis by ubiquitination and proteasome degradation has been investigated before^61,62^, however, as far as we know, there is no literature describing in which specific sites PTK2 is ubiquitinated or confirming its proteolytic degradation by proteasomes. Our co-immunoprecipitation assay coupled to mass spectrometry assay identified K926 as a target for ubiquitin-ligases (Fig. 8), a complex that recognizes and transfers a ubiquitin chain to lysine residues exposed in the surface of proteins containing degradations signals^63,64^. Next, we confirmed the UPS action on PTK2 homeostasis performing the pharmacological inhibition of proteasomes. The accumulation of ubiquitinated PTK2 in the nucleus, as revealed by super-resolution microscopy, provided strong evidence for the ubiquitin-proteasome system as a key regulator of nuclear PTK2 degradation (Fig. 7). Our findings on PTK2 quality control suggest a dynamic mobility of PTK2 within Bclaf1 biomolecular condensates, where this kinase is protected from UPS-mediated degradation. However, PTK2 diffused throughout the nucleus can be destabilized by oxidative damage, leading to its ubiquitination and proteasome degradation or to its accumulation in amorphous nuclear aggregates after an extended exposition to OS.

PTK2 regulates distinct cellular functions in cardiomyocytes, including survival during stress conditions^1,2,4,39^. Here, we present strong evidence that Bclaf1 condensates protect PTK2 against the UPS in a crucial mechanism for enhanced cardiomyocyte survival during OS. Interestingly, Bclaf1 knockdown resulted in increased general protein ubiquitination, pointing towards a broad role for Bclaf1 in counteracting the UPS on proteostasis control (Fig. 9)^12,65^. Furthermore, we observed reduced PTK2 levels and an accumulation of p53 following Bclaf1 knockdown, ultimately leading to cell death via the activation of apoptotic pathways (Fig. 9). It is plausible that the reduction of PTK2 availability contributes to p53 accumulation and subsequent apoptosis activation. Previous studies have demonstrated a role of αβ-crystallin to protect PTK2 from cytoplasmic calpain mediated-degradation in cardiomyocytes under mechanical stress, allowing cells to survive^4^. Here, we present evidence that Bclaf1 condensates play a similar role stabilizing nuclear PTK2 during OS. Indeed, past research demonstrated Bclaf1 knockout sensitized cells to TNF induced apoptosis, suggesting a regulatory role for this protein on cell fate^66^.

Our findings strongly support the role of Bclaf1 biomolecular condensates in counteracting PTK2 degradation by the ubiquitin-proteasome system, thereby regulating PTK2 homeostasis to promote cardiomyocyte survival during doxorubicin-induced oxidative stress and cytotoxicity.

## 4. MATERIAL AND METHODS

### 4.1 Cell culture

H9c2 cardiomyocytes, derived from rat embryos ventricular myoblasts, were obtained from American Type Culture Collection (ATCC). Cells were cultured in high-glucose DMEM supplemented with 3.7 g/L sodium bicarbonate, 10% FBS, and 1% penicillin/streptomycin solution (Sigma-Aldrich, P0781) at 37 °C with 5% CO_2_ atmosphere in a humidified incubator. Medium was changed every 2 days, and cells were trypsinized (Trypsin-EDTA) and split as soon as they reached around 70% confluence. Cells were counted using a Neubauer chamber to determine the appropriate seeding densities for each experiment.

### 4.2 Cell treatments

#### Doxorubicin

Doxorubicin powder was dissolved on water at 2 mg/mL. Then, for H9c2 cardiomyocytes treatments, dox was diluted on the complete medium (DMEM) with FBS and antibiotic in a final concentration of 1 µM. For recovery experiments, cells were treated with dox (1 µM) for 3 or 12 hours and then were incubated in complete medium for 12 and 36 hours. The other experiments were performed after 12 hours of dox treatment.

#### MG132

MG132 proteasome inhibitor (Sigma, M7449) was diluted on complete DMEM medium at 5 µM and cells were treated, in combination with dox (1 µM), for 12 hours. Control cells were treated with DMSO (vehicle) at the same volume as the proteasome inhibitor. After 12 hours, cells were fixed, or samples were extracted.

#### VER-155008

VER-155008 Hsp70 inhibitor (Sigma, SML0271) was diluted with DMSO as a stock solution at 5 mM and used at a final concentration of 50 µM^67^. Cells were treated for 3 h with dox (1 µM) followed by 1h of the Hsp70 inhibitor. For the experiments, the medium was changed to DMEM with FBS and antibiotics.

### 4.3 Coimmunoprecipitation assays

Proteins were extracted from H9c2 cells using Pierce™ Classic IP Kit (Thermo Fisher, 26146) with lysis buffer and protease inhibitor cocktail with EDTA on ice. Cells were lysed for 5 minutes using a cell scraper. Each sample was collected, agitated on a vortex, and stored in ice for 15 min. Next, samples were centrifuged to 12000 RPM for 20 minutes at 4°C. The pellet was discarded, and the supernatant was stored at -20°C for the co-immunoprecipitation assays. Protein quantification was performed using the BCA Protein Assay Kit (Thermo Fisher, 23225). Equal amounts of total extracts were used for co-immunoprecipitation using specific antibodies against PTK2 and Bclaf1 according to the manufacturer’s instruction.

### 4.4 Western blotting

H9c2 cells were lysed for 5 minutes using a RIPA buffer on ice, using a cell scraper. Each sample was collected, agitated on a vortex, and stored in ice for 15 min. Then, samples were centrifuged to 12000 RPM for 20 minutes at 4°C. The pellet was discarded, and the supernatant was quantified using Pierce BSA Assay Kit (Thermo Fisher, A55864) and then diluted on a 2X Laemmli loading buffer with β-mercaptoethanol. Next, samples were separated on 6%, 8%, or 12% SDS-polyacrylamide gels using prestained PageRuler (SM0761, Fermentas) as protein ladder. Proteins were transferred from gels to PVDF membranes, which were blocked with 5% BSA-TBS-T solution under agitation for 40 minutes. Then, the membranes were washed with TBS-T once and incubated with primary antibodies against PTK2 (Thermo Fisher, AHO0502), pPTK2 (Thermo Fisher, GTX50185), Bclaf1 (Thermo Fisher, PA578299), p53 (Thermo Fisher, MA1-7629), p-p53 (Thermo Fisher, PA5-117179), cleaved caspase-3 (Thermo Fisher, PA5114687), ubiquitin (Thermo Fisher, 13-1600), PUMA (Thermo Fisher, MA531994), GAPDH (SC-25778), α-tubulin (Sigma, T9026) and histone H1 (Thermo Fisher, PA5-30055) overnight. The dilution used for primary antibodies was 1µg/mL in 3% BSA-TBS-T solution. Next, the membranes were washed 3 times (10 minutes) with TBS-T and incubated with peroxidase conjugated secondary antibodies Goat Anti-Rabbit IgG (H + L)-HRP Conjugate (Bio-Rad, 1706515) or Goat Anti-Mouse IgG (H + L)-HRP Conjugate (Bio-Rad, 1706516) in 3% BSA-TBS-T solution for 1 hour and 30 minutes under mild agitation. After the incubation, membranes were washed 3 times (10 minutes) with TBS-T and visualized using the Clarity Western ECL Blotting Substrate (Bio-Rad, 1705060) in a Uvitec Documentation System.

### 4.5 Nuclear and Cytoplasmic Extraction

H9c2 cells were trypsinized and the nuclear and cytoplasmic protein extracts were obtained using NE-PER Nuclear and Cytoplasmic Extraction Reagents (Thermo Fisher, 78833), following the fabricant instructions.

### 4.6 Soluble and insoluble extraction

Proteins were extracted from H9c2 cells using Tris-saline solution (Tris-HCl: 50 mmol/L, NaCl: 150 mmol/L; pH: 7,6) containing protease inhibitor cocktail with EDTA on ice. Cells were scraped off the plates using a cell scraper, followed by sonication for 15 seconds, repeated for three times. The samples were then centrifuged at 12000 RPM for 30 minutes at 4°C and the supernatant was separated from the pellet. The pellet was solubilized in a solution of urea 6 M; SDS 2%; Tris-base 50 mM (pH 7,5); DTT 50 mM and incubated in a dry bath for 15 min at 60°C. Next, 5X Laemmli loading buffer with β-mercaptoethanol was added to all samples, followed by incubation in a dry bath for 5 minutes at 95°C. Finally, the samples were stored at -20°C.

### 4.7 Immunofluorescence

H9c2 cells were counted and seeded on 24 well plates containing coverslip (ThorLabs, 0117520). An amount of 2x10^4^ cells were plated for well, which were maintained in high glucose DMEM supplemented with 3.7 g/L sodium bicarbonate, 10% FBS and 1% penicillin/streptomycin solution. After cell adhesion to the coverslip, the treatments were conducted as described above. Then, cells were fixed using 4% paraformaldehyde, blocked and permeabilized using a solution with 3% bovine serum albumin and 0.1 % Triton-X in 0.1 mol/L DPBS on ice. Samples were incubated with primary antibodies against Bclaf1 (Thermo Fisher, PA5-78299 - 1:200 or Thermo Fisher, PA5-78299 - 1:200) and ubiquitin (Thermo Fisher, 13-600 - 1:100) for 30 minutes at room temperature. Other combinations were performed with antibodies against phosphorylated PTK2 (Thermo Fisher, 44-625G - 1:200) and Hsp70 (Thermo Fisher, MA3-006 - 1:50). Next, after rinsing, cells were labeled with Alexa Fluor-546-conjugated goat anti-rabbit (Thermo Fisher, A-11010 - 1:2000), Alexa Fluor-647-conjugated goat anti-mouse (Thermo Fisher, A-21235 - 1:2000) for 30 minutes at room temperature. Then, cells were labeled using FAK Monoclonal Antibody Alexa Fluor-488 (Thermo Fisher, MA3-96500-A488 - 1:200) for 30 minutes at room temperature. When the actin cytoskeleton was stained, Phalloidin-Alexa Fluor 647 was used (Thermo Fisher, A22287 - 1:40). After each incubation, the excess of antibodies was removed, washing 3 times using 1 mL of DPBS per well and agitating the plate on an orbital shaker for 5 minutes. Slides were mounted using ProLong Gold Antifade Mountant with DAPI.

### 4.8 Proximity Ligation Assay

H9c2 cells were plated on glass slides and fixed with 10% buffered formalin. Next, samples were incubated with glycine 100 mM and blocked using Duolink Blocking Solution (Sigma, DUO82007). Then, primary antibodies against PTK2 (Thermo Fisher, ZF002, mouse) and Bclaf1 (Thermo Fisher, PA5-78299, rabbit) were added to a 1:200 concentration and incubated in a humidity chamber at 4°C for 1 hour. The excess of antibodies was removed, and MINUS (DUO92004-100RXN, Sigma) and PLUS PLA (Sigma, DUO92002-100RXN) probes were added and incubated in a humidity chamber at 37°C for 1h and 10 minutes. Next, samples were incubated with ligation solution for 30 minutes at 37°C. Finally, an amplification buffer was added, and slides were incubated in a humidity chamber for 100 minutes at 37°C. After each step, samples were washed for 5 minutes at room temperature using a wash buffer. For negative controls, the samples were incubated with single primary antibodies or only the secondary antibodies and processed as in the PLA assay.

### 4.9 CellROX

H9c2 cells were plated on glass slides and treated with dox at 1 µM concentration during different times (1h, 3h, 6h and 12h). Next, CellROX Green Reagent (Thermo Fisher, C10444) was added at 5 μM concentration and incubated at 37°C for 30 minutes on an incubator. Then, cells were fixed using 4% paraformaldehyde solution and images were obtained in a confocal microscope using a 488 nm laser.

### 4.10 Transfection

For the Fluorescence lifetime imaging microscopy (FLIM) and Fluorescence recovery after photobleaching (FRAP) assays, cells were plated in a 35 mm glass bottom dish with 20 mm microwell (Cellvis, D35C4-20-0-N) and transfected with lipofectamine reagent (Invitrogen, L3000015) following the manufacturer’s instructions, using a medium supplemented with 5% FBS. After 1 day of transfection, the medium was replaced by a supplemented medium with 10% FBS for 24 hours. Then, the cells were treated with dox at 1 µM concentration during 3 hours. For experiments with Hsp70 inhibitor, cells were treated for 1 hour with VER-155008 after the treatment with dox. Cells were transfected with the EGFP-PTK2 or with the EGFP-Bclaf1 vectors (SynBio Technologies).

### 4.11 FLIM and FRAP

Experiments were performed on a LSM780-NLO Microscope (Carl Zeiss AG, Germany) using the EC Plan-Neofluar 40x/1.30 Oil DIC objective. Cells were maintained at 37 C° and 5% CO2 during acquisition. FLIM images were acquired using a TCSPC system from Becker-Hickl - Germany with a Hybrid-PMT HPM-100-40 detector. GFP fluorescence was excited at 488 nm by Two-Photon Excited Fluorescence using a ChameleonDiscovery NX (Coherent - USA) operating at 976 nm. In the FRAP experiments, a time series of 200 images were taken with a speed of 9 frames per second up to 5 min with an imaging area of 512 pixels × 512 pixels. The regions of interest were bleached down to 80% of original fluorescence signal excited by the 488 nm laser of the microscope. Curves were normalized by 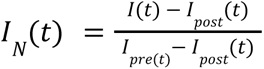, where *I_post_* (*t*) is the fluorescence intensity of the first frame post bleaching and *I_pre_* (*t*) is the fluorescence intensity of the last pre bleaching frame, this way all the curves were normalized to have pre bleaching intensity of 1 and post bleaching intensity of 0. Plotted curves are the average signal of the experiments and error bars are the standard deviation.

### 4.12 MTT

H9c2 cells were seeded at 1 x 10^4^ per well in a 96 well plate, for the MTT assays. After adhesion, cells were treated with dox during distinct times (1h, 3h, 6h and 12h). Then, the medium was replaced for the MTT solution (0.5 mg/mL), and cells were maintained for 4 hours at 37°C in an incubator. After the incubation, the precipitated crystals were diluted in DMSO and incubated for 10 minutes at 37°C. Then, in a plate reader, the absorbance was measured on a wavelength of 540 nm to determine cell viability. Seven replicates were done.

### 4.13 Short Hairpin RNA-mediated gene silencing

The short hairpin RNA (shRNA) sequence targeting Bclaf1 (*Rattus norvegicus*) was retrieved from the Broad Institute portal (https://portals.broadinstitute.org/). We selected a sequence located after 150 bp of the beginning of the coding sequence (CDS) with a GC content between 25% and 60% and excluding any candidate with a run of 7 C/G bases. The shRNA selected (5’-TGCTACATCTGGTGATATTTG-3’) was cloned into the plasmid PlKO.1 hygro (Addgene, #24150) previously digested with AgeI (NEB, #R0552S) and EcoRI (NEB, #R0101S) enzymes. The cloned plasmid was co-transfected with pCMV delta R8.2 (Addgene, #12263), and pMD2.G (Addgene, #12259) into Lenti-X 293T Cell Line (Takara Bio) for lentivirus packaging. The viral supernatant was then used to transduce H9c2 cells in the presence of polybrene (8 μg/mL) (Sigma-Aldrich, H9268) for 18 hours. Five days post-transduction, the selection was started using hygromycin (Thermo Fisher, Gibco 10687010) at a dose of 200 μg/mL for 5 days.

### 4.14 Microscopy analysis

SR-SIM images were acquired on a Zeiss Elyra PS.1 microscope, using PlanApochromat 63x/1.4 Oil DIC objective in the 3D-SIM mode. 3D-SIM z-stacks were projected on a single plane with summed intensities using the ImageJ-FIJI software. Correlation analysis was performed following the same method described by Galiani *et al*.^68^, using Bclaf1 clusters as origins to draw circles and calculate the integrated density fluorescence correlation. Chromatic 3D correction was performed by imaging different z planes of fluorescent sub-resolution beads using the same filter configuration for each channel as on the cell images. The resulting image was affine transformed by Zen Software (Zeiss) using the 488 nm channel as base position.

### 4.15 RT-qPCR

Total RNA was extracted using TRIzol according to Chomczynski & Sacchi^69^. One microgram was reverse transcribed using random primers and the Omniscript RT Kit (Qiagen, 205113) polymerase kit. Primers were designed using Primer3 software (primer3plus.com). The primer concentrations used in qPCR reactions were defined by testing different concentrations (200 nM, 400 nM and 800 nM). The lowest concentration without impact on product amplification and dimer formation was used for the experiments. The GAPDH gene was used as an endogenous control. The primers sequences are as follows: PTK2—5’GAGACCATTCCCATCCTTCCA3’ and 5’GTCGGAGTTCAGCAGCTTCT3’ (200 nM). GAPDH—5’GGAGCGAGATCCCTCCAAAAT3’ and 5’GGCTGTTGTCATACTTCT CATGG3’ (200 nM). For qPCR reactions, we used SYBR Green PCR Kit (Quantinova, 208052), 50 ng of cDNA, primers (200 nM) and Milli-Q water to reach the final reaction volume. Samples were assayed in triplicate and analysis was performed with a 7500 RT-PCR system (Applied Biosystems, California, USA).

### 4.16 Mass Spectrometry analysis of ubiquitinated sites

Following the PTK2 immunoprecipitation, we performed on-bead protein digestion. Briefly, matrix-associated proteins were resuspended in ammonium bicarbonate (AmBic 50 mM) and digested using trypsin (1:50 enzyme-to-substrate ratio) overnight at 36°C. Peptides were acidified with 0.1% formic acid and 2 µg of the mixture was loaded onto an Ultimate 3000 system (Thermo Scientific) coupled to an Orbitrap Exploris 240 (Thermo Scientific). Peptides were separated using an Acclaim PepMap RSLC C18 column (Thermo Scientific) with a binary gradient of acetonitrile (ACN) in 0.1% formic acid (FA). The gradient increased from 1% to 40% ACN over 120 minutes. For mass spectrometry analysis, data-dependent acquisition (DDA) was employed, with Precursor ions being analyzed with a resolution of 60,000. The top 20 most intense ions with charges of 2+ and 3+ were selected for MS/MS fragmentation using normalized collision energy (NCE) of 25, 30, and 35. Additionally, MS/MS spectra were acquired at a resolution of 15,000 with a dynamic exclusion time of 10 seconds. LC-MS raw data were analyzed using PatternLab V software^70^. Peptide spectral match was carried out against the *Rattus norvegicus* protein database from UniProt (version: 2023_02). Search parameters included -GG variable modification (m/z = 114.042927) in lysines and n-terminus to search ubiquitination sites.

### 4.17 Structural predictions

Protein sequences were obtained from UniProt, and their structures and potential interaction models were predicted using AlphaFold-Multimer^24,25,25^. This specialized version of AlphaFold was used to predict both individual protein structures and protein complexes models. The accuracy of the predicted complex structures was assessed using the pTM score, with scores above 0.5 indicating potential similarity to the true structures. However, pTM scores should be interpreted cautiously, as they can be influenced by the accuracy of individual protein components. To visualize and analyze the predicted structures, we employed UCSF ChimeraX, which allowed us to create 3D models and align different AlphaFold-predicted models to select the most representative structure^24^. To evaluate the propensity of the proteins to form biomolecular condensates, we utilized liquid-liquid phase separation (LLPS) prediction software including FuzDrop and PSPredictor. The FuzDrop server has four main application areas related to condensed states: (i) proteins are analyzed for liquid-liquid phase separation (LLPS) by calculating their droplet formation probability (pLLPS), with a threshold of 0.60 indicating a high likelihood of driving droplet formation; (ii) protein regions that promote droplet formation are identified, focusing on those with a droplet-promoting probability (pDP) ≥ 0.60 and at least 10 residues; (iii) droplet-client proteins, which cannot spontaneously undergo LLPS but contain droplet-promoting regions, are also identified, as they can partition into droplets through interactions with other protein; (iv) and aggregation hot-spots within droplets are detected by combining predictions on droplet-promoting regions with interaction diversity data using FuzPred^30^. Additionally, we used PSPredictor, a machine learning-based tool that predicts propensity of proteins undergoing phase separation. LLPS propensity scores above 0.60 were considered indicative of a high probability for condensate formation. PSPredictor employs sophisticated algorithms and a curated database of phase-separating proteins to provide accurate LLPS predictions^71^.

### 4.18 Statistical analysis

Statistical analysis was performed on GraphPad Prism software using Shapiro-Wilk and Kolmogorov-Smirnov tests for normality of the sample distribution and ANOVA or Mann-Whitney U tests for the analysis of variance between samples. P values < 0.05 were considered statistically significant, while * means P ≤ 0.05, ** means P ≤ 0.01, *** means P ≤ 0.001 and **** means P ≤ 0.0001.

## AUTHOR CONTRIBUTIONS

Conceptualization and methodology: AMS, AAT, IAM, and BRIR. Cell culture, treatments and sample extraction: IAM, BRIR, AGA, APS, PVCC, MGSC, and AMS; Differentiation assays: LBN and AMS. Super-resolution microscopy experiments and data analysis: IAM, BRIR, MCS, IRB, LBN, AMS and AAT; FRAP and FLIM: BRIR, IAM, AMS and AAT; PLA assays, imaging and data analysis: FLB, IAM and JK; RT-qPCR: IAM and MVG; PTK2 primers design: MVG; Mass spectrometry and data analysis: GLA, GRO, PCC, MDMS, RD, AGA, IAM, AMS and FCG; Short hairpin RNA sequences design: CHGS; Short hairpin RNA-mediated gene knockdown: FVR and CHGS; Biological interpretation: IAM, BRIR, HFC, AAT, and AMS; Writing: IAM, BRIR, HFC, AAT, and AMS. All authors edited and revised the final version of the manuscript.

## ACKNOWLEDGEMENTS

We thank the access to equipment and assistance provided by the National Institute of Science and Technology on Photonics Applied to Cell Biology (INFABIC) at the State University of Campinas; INFABIC is co-funded by Fundação de Amparo à Pesquisa do Estado de São Paulo (FAPESP) (2014/50938-8) and Conselho Nacional de Desenvolvimento Científico e Tecnológico (CNPq) (465699/2014-6). This research was supported by the Coordenação de Aperfeiçoamento de Pessoal de Nível Superior – Brasil (CAPES) (88887.513417/2020-00), the National Council for Scientific and Technological Development (CNPq) (423228/2018-8), the São Paulo Research Foundation (FAPESP) (2021/02303-7, 2018/07383-6, and 2020/11824-8) and the Fundação de Desenvolvimento da UNICAMP (FUNCAMP) (87/23-2545/23).

## DATA AVAILABILITY

Proteomics data were deposited in the ProteomeXchange Consortium via the PRIDE partner repository (accession no. PXD057681).

## REFERENCES

1. Frame, M. C., Patel, H., Serrels, B., Lietha, D. & Eck, M. J. The FERM domain: organizing the structure and function of FAK. Nat. Rev. Mol. Cell Biol. 11, 802–814 (2010).

2. Santos, A. M. et al. FERM domain interaction with myosin negatively regulates FAK in cardiomyocyte hypertrophy. Nat. Chem. Biol. 8, 102–110 (2012).

3. Fonseca, P. M. et al. Targeting to C-Terminal Myosin Heavy Chain May Explain Mechanotransduction Involving Focal Adhesion Kinase in Cardiac Myocytes. Circ. Res. 96, 73–81 (2005).

4. Pereira, M. B. et al. αB-crystallin interacts with and prevents stress-activated proteolysis of focal adhesion kinase by calpain in cardiomyocytes. Nat. Commun. 5, 5159 (2014).

5. Mitra, S. K., Hanson, D. A. & Schlaepfer, D. D. Focal adhesion kinase: in command and control of cell motility. Nat. Rev. Mol. Cell Biol. 6, 56–68 (2005).

6. Zhou, J., Yi, Q. & Tang, L. The roles of nuclear focal adhesion kinase (FAK) on Cancer: a focused review. J. Exp. Clin. Cancer Res. CR 38, 250 (2019).

7. Schlaepfer, D. D., Hauck, C. R. & Sieg, D. J. Signaling through focal adhesion kinase. Prog. Biophys. Mol. Biol. 71, 435–478 (1999).

8. Marin, T. M. et al. Shp2 negatively regulates growth in cardiomyocytes by controlling focal adhesion kinase/Src and mTOR pathways. Circ. Res. 103, 813–824 (2008).

9. Senyo, S. E., Koshman, Y. E. & Russell, B. Stimulus interval, rate and direction differentially regulate phosphorylation for mechanotransduction in neonatal cardiac myocytes. FEBS Lett. 581, 4241–4247 (2007).

10. Nadruz, W., Corat, M. A. F., Marin, T. M., Guimarães Pereira, G. A. & Franchini, K. G. Focal adhesion kinase mediates MEF2 and c-Jun activation by stretch: role in the activation of the cardiac hypertrophic genetic program. Cardiovasc. Res. 68, 87–97 (2005).

11. Clemente, C. F. M. Z. et al. Focal adhesion kinase governs cardiac concentric hypertrophic growth by activating the AKT and mTOR pathways. J. Mol. Cell. Cardiol. 52, 493–501 (2012).

12. Lim, S.-T. et al. Nuclear FAK promotes cell proliferation and survival through FERM-enhanced p53 degradation. Mol. Cell 29, 9–22 (2008).

13. Cheng, Z. et al. Focal adhesion kinase antagonizes doxorubicin cardiotoxicity via p21(Cip1.). J. Mol. Cell. Cardiol. 67, 1–11 (2014).

14. Tang, K.-J. et al. Focal Adhesion Kinase Regulates the DNA Damage Response and Its Inhibition Radiosensitizes Mutant KRAS Lung Cancer. Clin. Cancer Res. Off. J. Am. Assoc. Cancer Res. 22, 5851–5863 (2016).

15. Tavora, B. et al. Endothelial-cell FAK targeting sensitizes tumours to DNA-damaging therapy. Nature 514, 112–116 (2014).

16. Ma, Z. et al. Role of BCLAF-1 in PD-L1 stabilization in response to ionizing irradiation. Cancer Sci. 112, 4064–4074 (2021).

17. Shao, A. et al. Bclaf1 is a direct target of HIF-1 and critically regulates the stability of HIF-1α under hypoxia. Oncogene 39, 2807–2818 (2020).

18. Ramanathan, A., Savol, A., Burger, V., Chennubhotla, C. S. & Agarwal, P. K. Protein conformational populations and functionally relevant substates. Acc. Chem. Res. 47, 149–156 (2014).

19. Ha, J.-H. & Loh, S. N. Protein conformational switches: from nature to design. Chem. Weinh. Bergstr. Ger. 18, 7984–7999 (2012).

20. Forman-Kay, J. D., Ditlev, J. A., Nosella, M. L. & Lee, H. O. What are the distinguishing features and size requirements of biomolecular condensates and their implications for RNA-containing condensates? RNA N. Y. N 28, 36–47 (2022).

21. Lietha, D. et al. Structural basis for the autoinhibition of focal adhesion kinase. Cell 129, 1177–1187 (2007).

22. Sanders, D. W. et al. Competing Protein-RNA Interaction Networks Control Multiphase Intracellular Organization. Cell 181, 306–324.e28 (2020).

23. Lin, Y., Protter, D. S. W., Rosen, M. K. & Parker, R. Formation and Maturation of Phase-Separated Liquid Droplets by RNA-Binding Proteins. Mol. Cell 60, 208–219 (2015).

24. Jumper, J. et al. Highly accurate protein structure prediction with AlphaFold. Nature 596, 583–589 (2021).

25. Abramson, J. et al. Accurate structure prediction of biomolecular interactions with AlphaFold 3. Nature 630, 493–500 (2024).

26. Yu, Z. et al. BCLAF1 binds SPOP to stabilize PD-L1 and promotes the development and immune escape of hepatocellular carcinoma. Cell. Mol. Life Sci. CMLS 81, 82 (2024).

27. Sarras, H., Alizadeh Azami, S. & McPherson, J. P. In search of a function for BCLAF1. ScientificWorldJournal 10, 1450–1461 (2010).

28. Alberti, S. & Hyman, A. A. Biomolecular condensates at the nexus of cellular stress, protein aggregation disease and ageing. Nat. Rev. Mol. Cell Biol. 22, 196–213 (2021).

29. Snead, W. T. & Gladfelter, A. S. The Control Centers of Biomolecular Phase Separation: How Membrane Surfaces, PTMs, and Active Processes Regulate Condensation. Mol. Cell 76, 295–305 (2019).

30. Hatos, A., Tosatto, S. C. E., Vendruscolo, M. & Fuxreiter, M. FuzDrop on AlphaFold: visualizing the sequence-dependent propensity of liquid-liquid phase separation and aggregation of proteins. Nucleic Acids Res. 50, W337–W344 (2022).

31. Liu, J. X. et al. Liquid-liquid phase separation within fibrillar networks. Nat. Commun. 14, 6085 (2023).

32. Alshareedah, I., Kaur, T. & Banerjee, P. R. Methods for characterizing the material properties of biomolecular condensates. Methods Enzymol. 646, 143–183 (2021).

33. Quan, M. D., Liao, S.-C. J., Ferreon, J. C. & Ferreon, A. C. M. Fluorescence Lifetime Imaging Microscopy of Biomolecular Condensates. Methods Mol. Biol. Clifton NJ 2563, 135–148 (2023).

34. Filomeni, G., De Zio, D. & Cecconi, F. Oxidative stress and autophagy: the clash between damage and metabolic needs. Cell Death Differ. 22, 377–388 (2015).

35. Scandalios, J. G. Oxidative stress: molecular perception and transduction of signals triggering antioxidant gene defenses. Braz. J. Med. Biol. Res. Rev. Bras. Pesqui. Medicas E Biol. 38, 995–1014 (2005).

36. Hirose, T., Ninomiya, K., Nakagawa, S. & Yamazaki, T. A guide to membraneless organelles and their various roles in gene regulation. Nat. Rev. Mol. Cell Biol. 24, 288–304 (2023).

37. Yoo, H., Bard, J. A. M., Pilipenko, E. V. & Drummond, D. A. Chaperones directly and efficiently disperse stress-triggered biomolecular condensates. Mol. Cell 82, 741–755.e11 (2022).

38. Hornbeck, P. V. et al. PhosphoSitePlus, 2014: mutations, PTMs and recalibrations. Nucleic Acids Res. 43, D512–520 (2015).

39. Sonoda, Y. et al. Anti-apoptotic role of focal adhesion kinase (FAK). Induction of inhibitor-of-apoptosis proteins and apoptosis suppression by the overexpression of FAK in a human leukemic cell line, HL-60. J. Biol. Chem. 275, 16309–16315 (2000).

40. Clemente, C. F. M. Z. et al. Targeting focal adhesion kinase with small interfering RNA prevents and reverses load-induced cardiac hypertrophy in mice. Circ. Res. 101, 1339–1348 (2007).

41. Focal adhesion kinase signaling in unexpected places - PubMed. https://pubmed.ncbi.nlm.nih.gov/28213315/.

42. Rodríguez-Fernández, J. L. Why do so many stimuli induce tyrosine phosphorylation of FAK? BioEssays News Rev. Mol. Cell. Dev. Biol. 21, 1069–1075 (1999).

43. Autophosphorylation of the focal adhesion kinase, pp125FAK, directs SH2-dependent binding of pp60src - PubMed. https://pubmed.ncbi.nlm.nih.gov/7509446/.

44. Liu, Y., Yao, Z., Lian, G. & Yang, P. Biomolecular phase separation in stress granule assembly and virus infection. Acta Biochim. Biophys. Sin. 55, 1099–1118 (2023).

45. Laflamme, G. & Mekhail, K. Biomolecular condensates as arbiters of biochemical reactions inside the nucleus. Commun. Biol. 3, 773 (2020).

46. Ranganathan, S., Liu, J. & Shakhnovich, E. Different states and the associated fates of biomolecular condensates. Essays Biochem. 66, 849–862 (2022).

47. Spegg, V. & Altmeyer, M. Biomolecular condensates at sites of DNA damage: More than just a phase. DNA Repair 106, 103179 (2021).

48. Wen, Y. et al. Bclaf1 promotes angiogenesis by regulating HIF-1α transcription in hepatocellular carcinoma. Oncogene 38, 1845–1859 (2019).

49. Shcherbakova, L., Pardo, M., Roumeliotis, T. & Choudhary, J. Identifying and characterising Thrap3, Bclaf1 and Erh interactions using cross-linking mass spectrometry. Wellcome Open Res. 6, 260 (2021).

50. Pliss, A. et al. Cycles of protein condensation and discharge in nuclear organelles studied by fluorescence lifetime imaging. Nat. Commun. 10, 455 (2019).

51. Case, L. B., De Pasquale, M., Henry, L. & Rosen, M. K. Synergistic phase separation of two pathways promotes integrin clustering and nascent adhesion formation. eLife 11, e72588 (2022).

52. Akerfelt, M., Morimoto, R. I. & Sistonen, L. Heat shock factors: integrators of cell stress, development and lifespan. Nat. Rev. Mol. Cell Biol. 11, 545–555 (2010).

53. Tyedmers, J., Mogk, A. & Bukau, B. Cellular strategies for controlling protein aggregation. Nat. Rev. Mol. Cell Biol. 11, 777–788 (2010).

54. Mitrea, D. M., Mittasch, M., Gomes, B. F., Klein, I. A. & Murcko, M. A. Modulating biomolecular condensates: a novel approach to drug discovery. Nat. Rev. Drug Discov. 21, 841–862 (2022).

55. Mateju, D. et al. An aberrant phase transition of stress granules triggered by misfolded protein and prevented by chaperone function. EMBO J. 36, 1669–1687 (2017).

56. Kong, C.-Y. et al. Underlying the Mechanisms of Doxorubicin-Induced Acute Cardiotoxicity: Oxidative Stress and Cell Death. Int. J. Biol. Sci. 18, 760–770 (2022).

57. Cappetta, D. et al. Oxidative Stress and Cellular Response to Doxorubicin: A Common Factor in the Complex Milieu of Anthracycline Cardiotoxicity. Oxid. Med. Cell. Longev. 2017, 1521020 (2017).

58. Zhang, X. et al. FNDC5 alleviates oxidative stress and cardiomyocyte apoptosis in doxorubicin-induced cardiotoxicity via activating AKT. Cell Death Differ. 27, 540–555 (2020).

59. Song, I.-K. et al. Degradation of Redox-Sensitive Proteins including Peroxiredoxins and DJ-1 is Promoted by Oxidation-induced Conformational Changes and Ubiquitination. Sci. Rep. 6, 34432 (2016).

60. Shang, F. & Taylor, A. Ubiquitin-proteasome pathway and cellular responses to oxidative stress. Free Radic. Biol. Med. 51, 5–16 (2011).

61. Fan, Y., Qu, X., Ma, Y., Liu, Y. & Hu, X. Cbl-b promotes cell detachment via ubiquitination of focal adhesion kinase. Oncol. Lett. 12, 1113–1118 (2016).

62. BommaReddy, R. R., Patel, R., Smalley, T. & Acevedo-Duncan, M. Effects of Atypical Protein Kinase C Inhibitor (DNDA) on Lung Cancer Proliferation and Migration by PKC-ι/FAK Ubiquitination Through the Cbl-b Pathway. OncoTargets Ther. 13, 1661–1676 (2020).

63. Nandi, D., Tahiliani, P., Kumar, A. & Chandu, D. The ubiquitin-proteasome system. J. Biosci. 31, 137–155 (2006).

64. Ranek, M. J. & Wang, X. Activation of the ubiquitin-proteasome system in doxorubicin cardiomyopathy. Curr. Hypertens. Rep. 11, 389–395 (2009).

65. Lee, Y. Y., Yu, Y. B., Gunawardena, H. P., Xie, L. & Chen, X. BCLAF1 is a radiation-induced H2AX-interacting partner involved in γH2AX-mediated regulation of apoptosis and DNA repair. Cell Death Dis. 3, e359 (2012).

66. Zhang, R. et al. Bclaf1 regulates c-FLIP expression and protects cells from TNF-induced apoptosis and tissue injury. EMBO Rep. 23, e52702 (2022).

67. Frottin, F. et al. The nucleolus functions as a phase-separated protein quality control compartment. Science 365, 342–347 (2019).

68. Galiani, S. et al. Super-resolution Microscopy Reveals Compartmentalization of Peroxisomal Membrane Proteins. J. Biol. Chem. 291, 16948–16962 (2016).

69. Chomczynski, P. & Sacchi, N. Single-step method of RNA isolation by acid guanidinium thiocyanate-phenol-chloroform extraction. Anal. Biochem. 162, 156–159 (1987).

70. Santos, M. D. M. et al. Simple, efficient and thorough shotgun proteomic analysis with PatternLab V. Nat. Protoc. 17, 1553–1578 (2022).

71. Sun, J. et al. Precise prediction of phase-separation key residues by machine learning. Nat. Commun. 15, 2662 (2024).

