## Supplemental Material for "Bclaf1 biomolecular condensates protect nuclear PTK2 from ubiquitin-proteasome system promoting cardiomyocyte survival during oxidative stress"

### Supplementary Results

**Supplementary Table 1.** Proteins identified in immunoprecipitated extracts.

| Gene Symbol | Entrez Gene Name | Sequence | XCorr | *m/z* (Da) | Charge |
|---|---|---|---|---|---|
| PTK2 / FAK | Protein tyrosine kinase 2 / Focal adhesion kinase | SEEVHWLHVDMGVSSVR | 4.68 | 983.97156 | 2 |
| | | YELAHPPEEWKYELR | 4.09 | 980.48560 | 2 |
| | | YELAHPPEEWKYELR | 3.82 | 653.98737 | 3 |
| | | YELAHPPEEWK | 3.78 | 699.83350 | 2 |
| | | QFANLNREESILK | 3.78 | 781.41711 | 2 |
| | | QFANLNREESILK | 3.76 | 781.41528 | 2 |
| | | YELAHPPEEWK | 3.72 | 699.83331 | 2 |
| | | YELAHPPEEWK | 3.66 | 699.83350 | 2 |
| | | EKYELAHPPEEWK | 3.48 | 828.40369 | 2 |
| | | EKYELAHPPEEWK | 3.40 | 828.40265 | 2 |
| | | YELAHPPEEWK | 3.30 | 699.83331 | 2 |
| | | FFEILSPVYR | 3.22 | 635.84259 | 2 |
| | | FFEILSPVYR | 3.11 | 635.84247 | 2 |
| | | EKYELAHPPEEWK | 3.00 | 828.40167 | 2 |
| | | EKYELAHPPEEWK | 2.98 | 828.40240 | 2 |
| | | SNYEVLEK | 2.90 | 491.24432 | 2 |
| | | SNYEVLEK | 2.84 | 491.24414 | 2 |
| | | KGMLQLK | 2.67 | 409.24841 | 2 |

| | | | | | |
|---|---|---|---|---|---|
| | | FFEILSPVYR | 2.66 | 635.84192 | 2 |
| | | KLIQQTFR | 2.61 | 517.30762 | 2 |
| | | IVDSHKVK | 2.53 | 463.27249 | 2 |
| | | SNYEVLEK | 2.47 | 491.24402 | 2 |
| | | GMLQLK | 2.33 | 345.20135 | 2 |
| | | KGMLQLK | 2.25 | 409.24875 | 2 |
| | | SNYEVLEK | 2.12 | 981.48059 | 1 |
| | | SNYEVLEK | 2.10 | 981.48022 | 1 |
| | | LGcLEIR | 2.05 | 860.46063 | 1 |
| | | SYWEMR | 1.96 | 871.37262 | 1 |
| Bclaf1 | BCL-associated transcription factor 1 | MAPVPLDDSNRPASLTK | 4.29 | 906.46552 | 2 |
| | | SSATSGDIWPGLSAYDNSPR | 4.16 | 1040.98108 | 2 |
| | | SQEEPKDTFEHDPSESIDEFNK | 3.94 | 870.04382 | 3 |
| | | MAPVPLDDSNRPASLTK | 3.68 | 906.46490 | 2 |
| | | MAPVPLDDSNRPASLTK | 3.31 | 906.46637 | 2 |
| | | GTFHDDRDDGVDYWAK | 3.25 | 948.90594 | 2 |
| | | SQEEPKDTFEHDPSESIDEFNK | 2.98 | 652.78479 | 4 |
| | | LKDLFDYSPPLHK | 2.86 | 524.94977 | 3 |
| | | SSFYPDGGDQETAK | 2.85 | 751.32166 | 2 |
| | | LLASTLVHSVK | 2.80 | 584.35480 | 2 |
| | | GTFHDDRDDGVDYWAK | 2.75 | 632.93915 | 3 |
| | | KSVLADQGK | 2.66 | 945.52942 | 1 |
| | | SSFYPDGGDQETAK | 2.59 | 751.32147 | 2 |

A

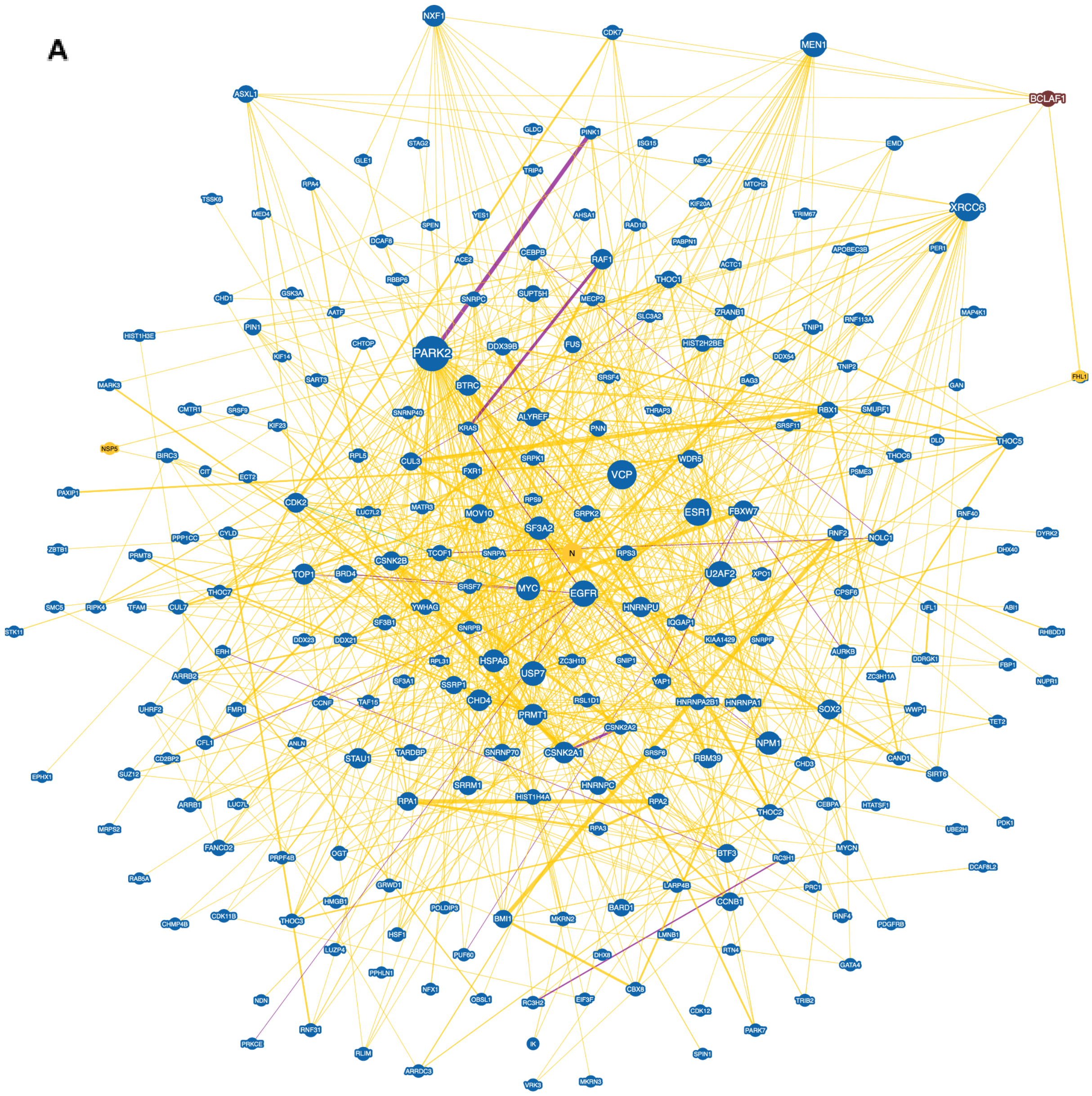
NXF1
CDK7
MEN1
BCLAF1
ASXL1
GLDC
PINK1
STAG2
ISG15
EMD
GLE1
NEK4
TRIP4
RPA4
MTCH2
TSSK6
KIF20A
XRCC6
MED4
YES1
AHSA1
TRIM67
SPEN
RAD18
DCAF8
PABPN1
PER1
APOBEC3B
CEBPB
ACE2
RAF1
ACTC1
RBBP6
THOC1
GSK3A
SUPT5H
CHD1
SNRPC
MECP2
ZRANB1
MAP4K1
AATF
SLC3A2
RNF113A
PIN1
TNIP1
HIST1H3E
CHTOP
DDX39B
FUS
HIST2H2BE
KIF14
PARK2
DDX54
TNIP2
MARK3
SART3
SRSF4
BAG3
FHL1
BTRC
GAN
CMTR1
SRSF9
SNRNP40
THRAP3
RBX1
SMURF1
KIF23
ALYREF
KRAS
PNN
SRSF11
DLD
THOC5
NSP5
BIRC3
RPL5
CUL3
SRPK1
WDR5
THOC6
CIT
FXR1
VCP
PSME3
PAXIP1
ECT2
CDK2
RPS9
LUC7L2
MATR3
MOV10
SRPK2
ESR1
FBXW7
RNF40
CYLD
SF3A2
RNF2
NOLC1
DYRK2
ZBTB1
PPP1CC
PRMT5
TCOF1
SNRPA
N
RPS3
DHX40
CSNK2B
BRD4
TOP1
U2AF2
XPO1
THOC7
SRSF7
MYC
EGFR
CPSF6
SMC5
RIPK4
TFAM
CUL7
YWHAG
HNRNPU
UFL1
ABI1
STK11
SF3B1
IQGAP1
RHBDD1
DDX23
DDX21
SNRPB
KIAA1429
SNRPF
ERH
AURKB
DDRGK1
RPL31
HSPA8
ZC3H18
SNIP1
FBP1
ARRB2
USP7
ZC3H11A
SF3A1
SSRP1
YAP1
NUPR1
UHRF2
FMR1
CCNF
TAF15
CHD4
RSL1D1
HNRNPA2B1
HNRNPA1
SOX2
WWP1
PRMT1
TET2
CFL1
ANLN
CSNK2A2
CD2BP2
TARDBP
SNRNP70
CSNK2A1
SRSF6
NPM1
STAU1
RBM39
CHD3
CAND1
EPHX1
SUZ12
SIRT6
SRRM1
HNRNPC
ARRB1
LUC7L
RPA1
HIST1H4A
RPA2
THOC2
CEBPA
HTATSF1
MRPS2
RPA3
UBE2H
PDK1
FANCD2
OGT
BTF3
MYCN
PRPF4B
RC3H1
RAB5A
DCAF8L2
GRWD1
LARP4B
PRC1
HMGB1
CCNB1
CDK11B
POLDIP3
BARD1
CHMP4B
THOC3
BMI1
MKRN2
RNF4
PDGFRB
HSF1
LMNB1
LUZP4
PUF60
DHX8
RTN4
GATA4
PPHLN1
CBX8
NFX1
NDN
OBSL1
RC3H2
EIF3F
CDK12
TRIB2
RNF31
PARK7
PRKCE
IK
RLIM
SPIN1
ARRDC3
MKRN3
VRK3

**B**

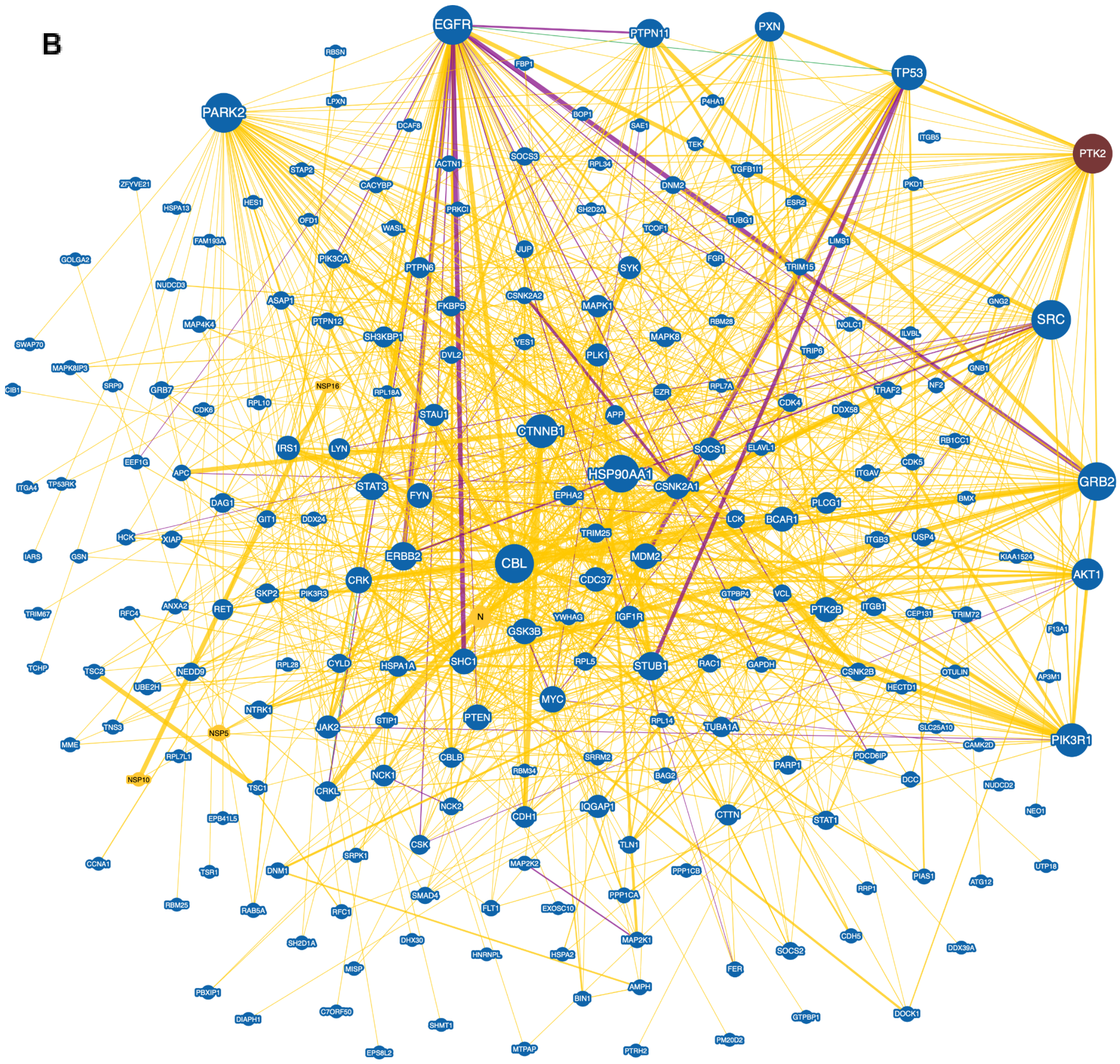


**Supplementary Figure 1. (A-B)** Bclaf1 and PTK2 protein-protein interaction network, respectively. The network was obtained from the Biogrid Platform (https://thebiogrid.org/), including physical and genetic interaction. The node size represents the enrichment of the pathway and the lines the degree of evidence. Some pathways, such as Park2, EGFR, and Myc, are enriched in both Bclaf1 and PTK2 pathways.

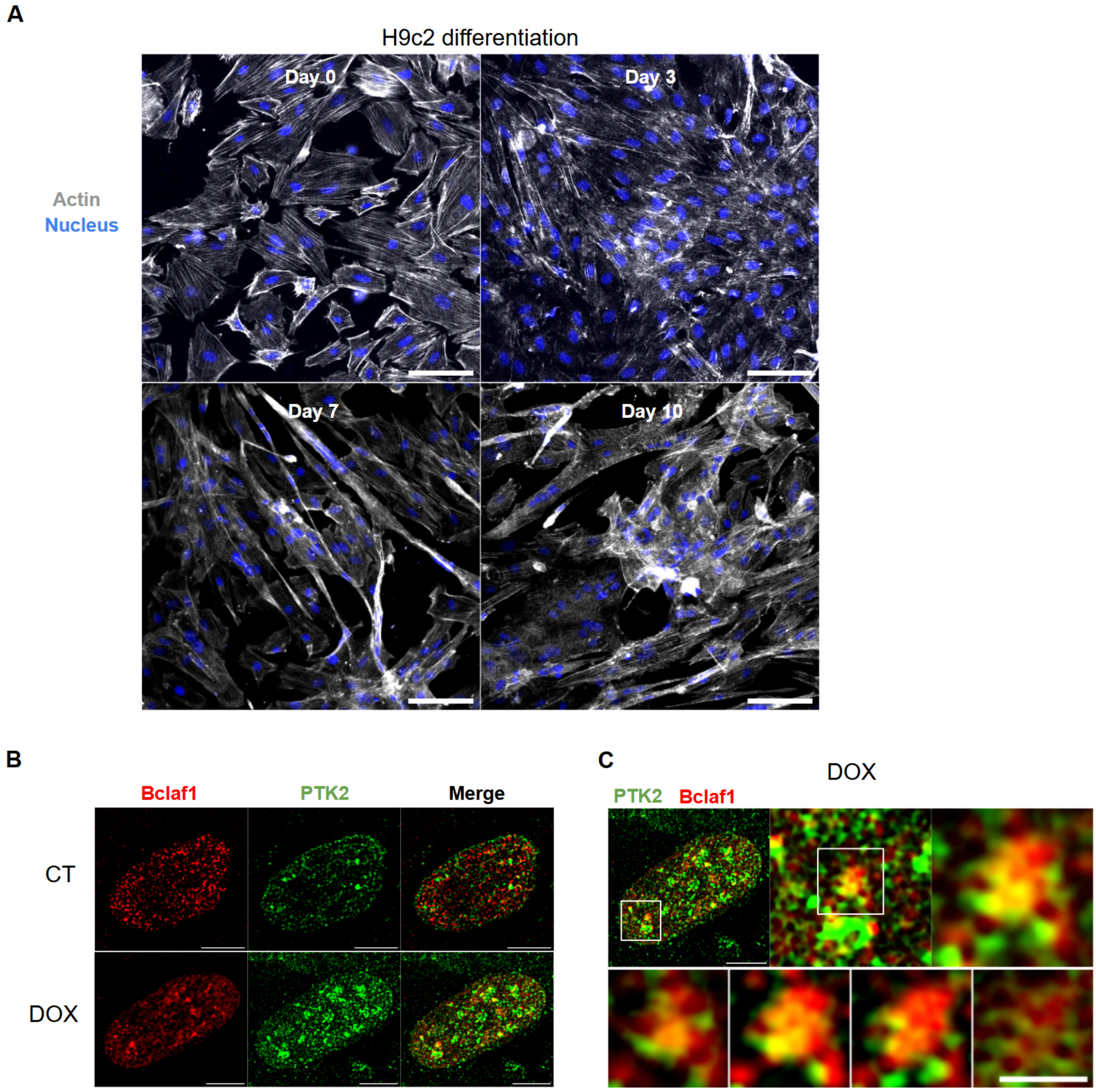


**Supplementary Figure 2. (A)** Immunofluorescence staining of actin and nuclei showing H9c2 cardiomyocyte differentiation over 10 days. Blue: nucleus; Gray: actin. Scale bar = 100 µm. **(B)** Immunofluorescence staining of Bclaf1 and PTK2 in differentiated H9c2 cardiomyocytes, showing subcellular localization of these proteins in control conditions and when cells were submitted to 12 hours of dox treatment (1 µM). Scale bar = 5 µm **(C)** Immunofluorescence staining of Bclaf1 and PTK2 in a differentiated H9c2 cardiomyocyte treated with dox, showing the colocalization of PTK2 and Bclaf1 in the nucleus. The bottom montage shows colocalization on nuclear condensates throughout 4 slices of the original three-dimensional image (Z-stack). Scale bar from Z-stack = 1 µm. Red: Bclaf1; Green: PTK2.

**A**

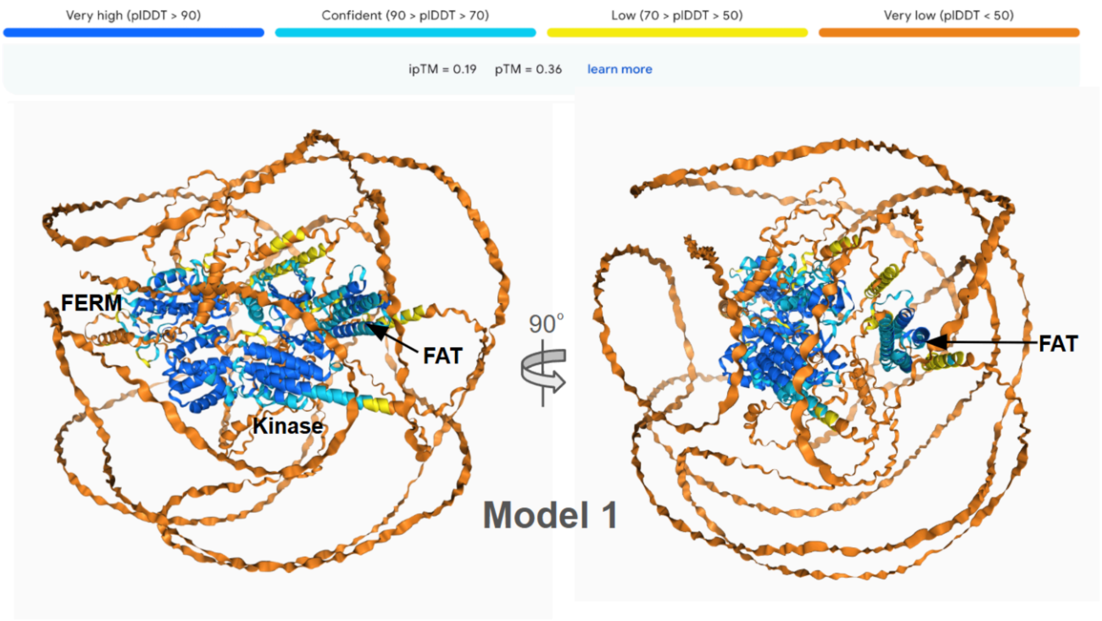


**B**

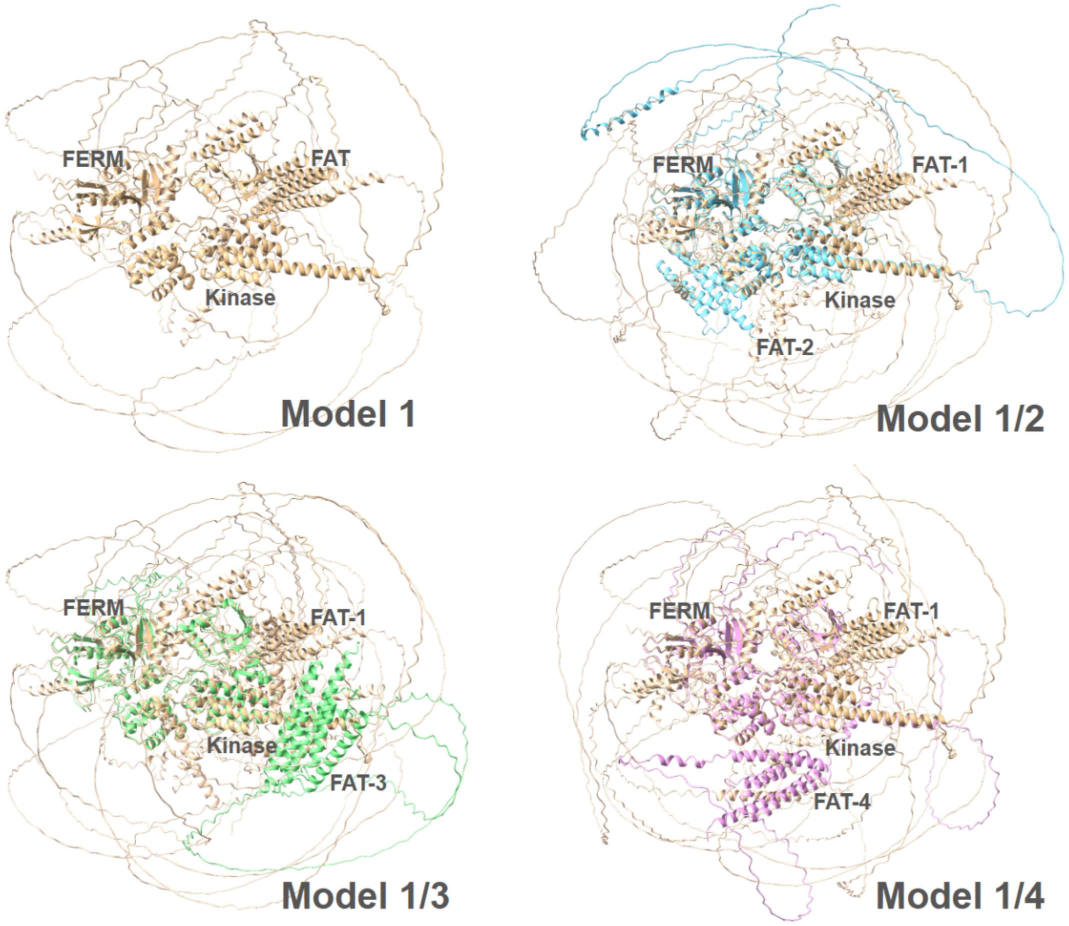


**Supplementary Figure 3.** Structural models of PTK2-Bclaf1 interaction. **(A)** Model 1 shows the primary structure of PTK2 with FERM (blue), kinase (green), and FAT (orange) domains.

The color coding represents confidence levels based on pLDDT scores. A 90° rotated view is shown. **(B)** Alignment of Model 1 with models 2, 3, and 4, highlighting the different positions of FAT domain in relation to FERM-Kinase (FAT-1, FAT-2, FAT-3, FAT-4), confirming the high flexibility provided by the linking between the kinase and FAT domains.

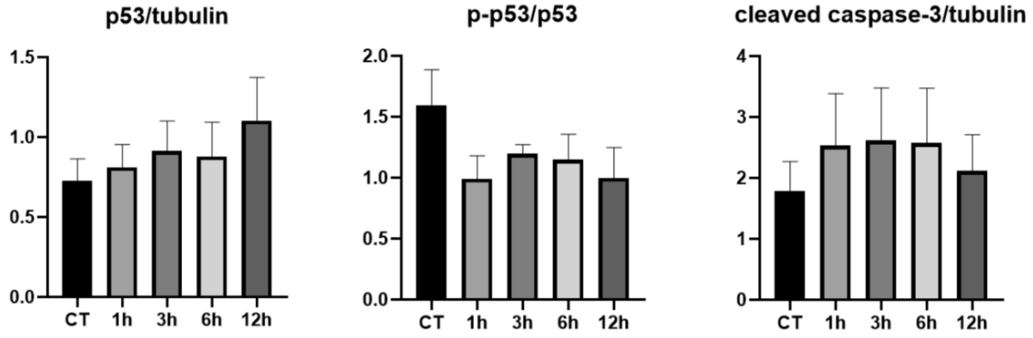


**Supplementary Figure 4.** Bar graph shows western blot analysis demonstrating phosphorylated p53 (p-p53), p53 and cleaved caspase-3, with no significant changes in expression. $\alpha$-Tubulin served as loading control.

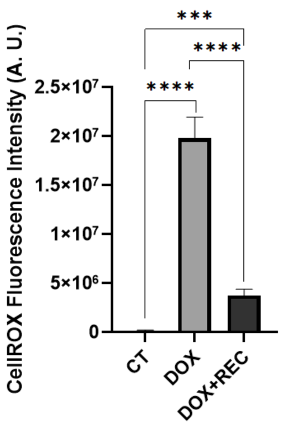


**Supplementary Figure 5.** Bar graphic of CellROX fluorescence after 36 hours of recovery (DOX + REC) from dox treatment (DOX). N = 7 cells per group.

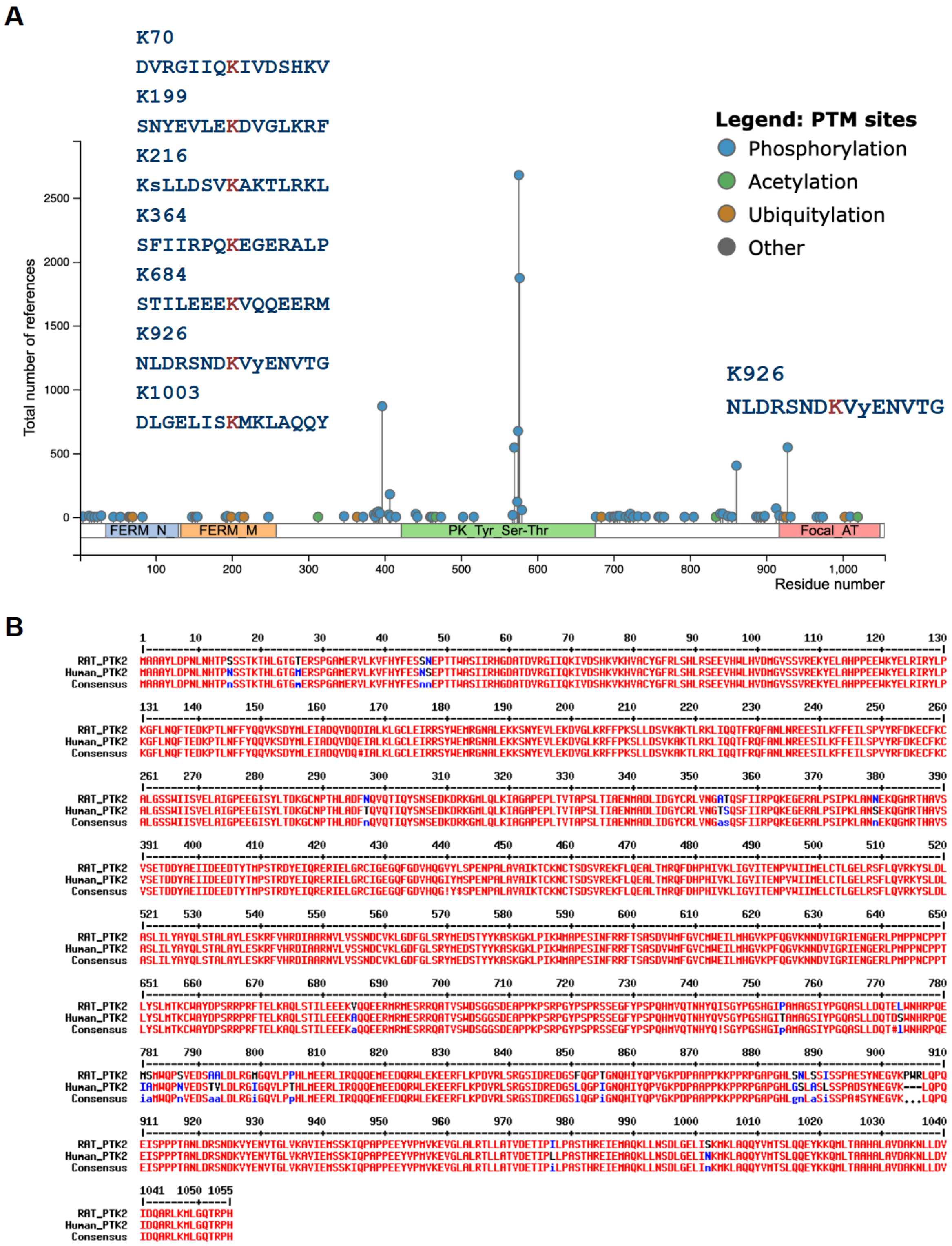


**Supplementary Figure 6. (A)** Rat PTK2 post-translational modification (PTM) mapping. PTM map showed probable phosphorylation (blue), acetylation (green), ubiquitylation (brown), and other modifications (gray) across the protein sequence. Key modified residues and their positions along the protein domains (FERM, kinase, and FAT) were highlighted. The seven probable ubiquitination sites on PTK2 are highlighted in the inset, including the peptide containing the K926 site. **(B)** Sequence alignment of PTK2 from rat (RAT_PTK2)

and human (Human_PTK2) obtained using MultAlin (http://multalin.toulouse.inra.fr/multalin/). The alignment shows the conserved regions between the two species, with identical residues highlighted in red. Differences between the sequences are marked in blue, while the consensus sequence is provided at the bottom for each alignment block. The numbers above indicate the position of the amino acid residues within the protein sequence. This alignment highlights both conserved and variable regions across species. The peptide sequence containing K926/K923 is conserved.

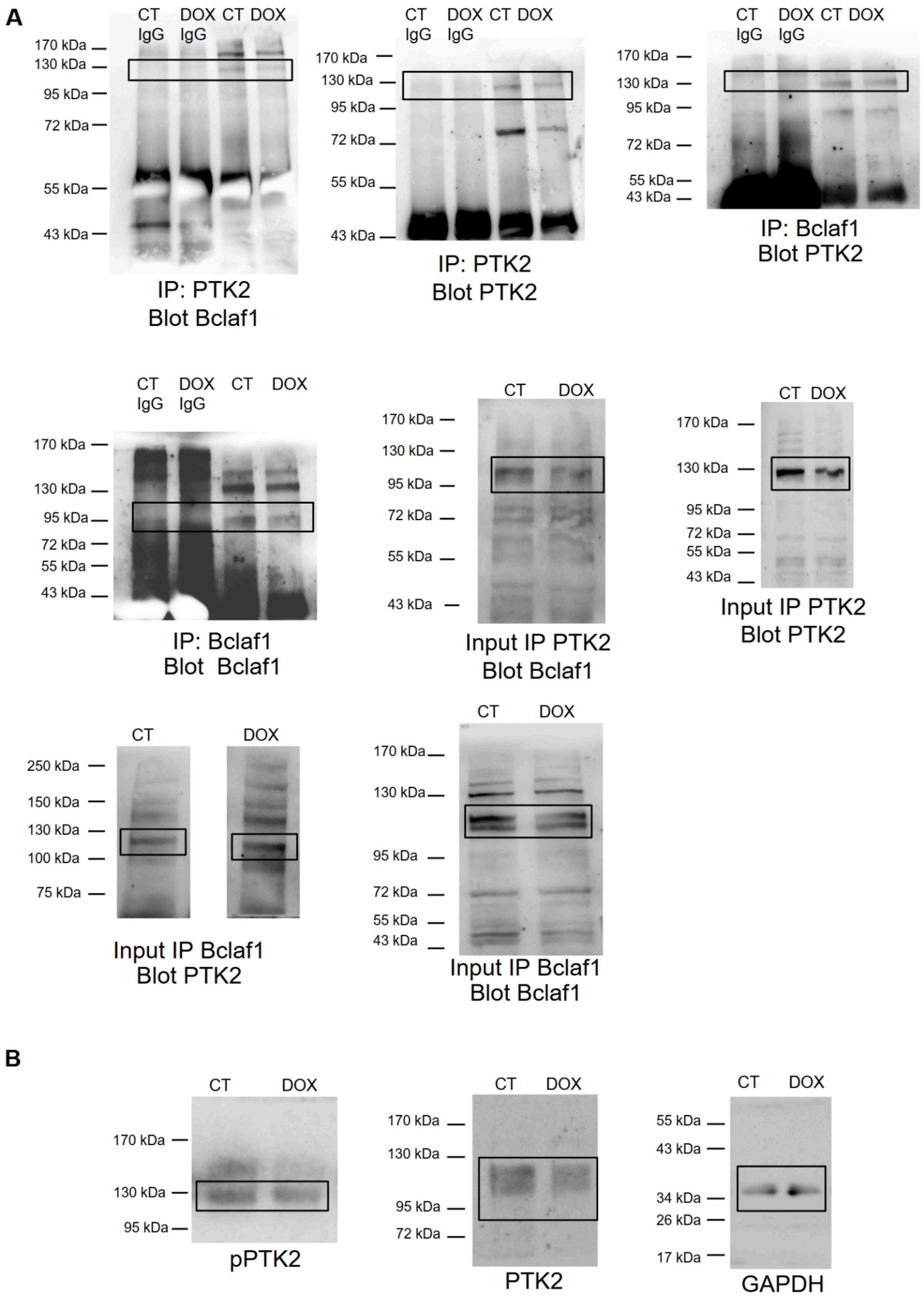
A
CT IgG
DOX IgG
CT
DOX
170 kDa
130 kDa
95 kDa
72 kDa
55 kDa
43 kDa
IP: PTK2
Blot Bclaf1
IP: PTK2
Blot PTK2
IP: Bclaf1
Blot PTK2
IP: Bclaf1
Blot Bclaf1
Input IP PTK2
Blot Bclaf1
Input IP PTK2
Blot PTK2
250 kDa
150 kDa
100 kDa
75 kDa
Input IP Bclaf1
Blot PTK2
Input IP Bclaf1
Blot Bclaf1
B
34 kDa
26 kDa
17 kDa
pPTK2
PTK2
GAPDH

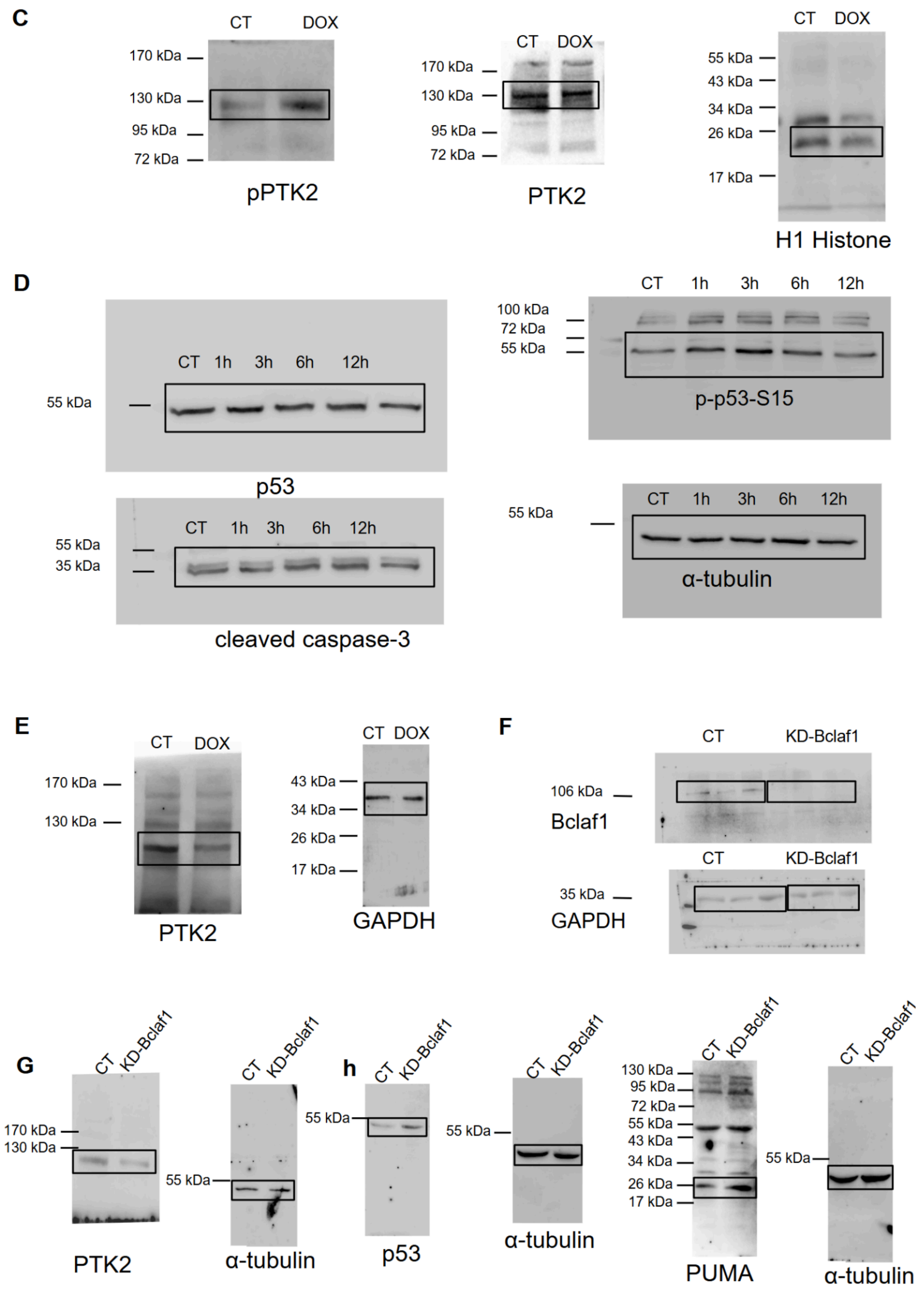


**Supplementary Figure 7.** Uncropped immunoblots. **(A)** Figure 1 and 3. **(B-C)** Figure 2. **(D)** Figure 6. **(E)** Figure 7. **(F-H)** Figure 9.

### Supplementary Methods

#### Co-Immunoprecipitation LC-MS/MS Analysis

HEK-293T control or expressing Flag-FERM were washed in cold PBS and scraped into lysis buffer (50 mM Tris-HCl pH 7.4, 150 mM NaCl, 1 mM EDTA, 1% Triton X-100, proteases and phosphatases inhibitors). The samples were centrifuged for 20 min at 16,000 g and equal amounts of protein (500 μg) were incubated with anti-FLAG M2 Affinity Gel (Sigma). Flag-immunocomplexes were eluted with a high concentration (150 ng/μL) of FLAG peptide (Sigma) during 2 hours at 4ºC. Subsequently, the eluted proteins were prepared by mass spectrometry.

For a bottom-up study, Flag-FERM-PTK2 and the interacting partners were reduced with dithiothreitol (5 mM for 25 min at 56 °C) and alkylated with iodoacetamide (14 mM for 30 min at room temperature in the dark). The samples were digested with trypsin (Promega) and dried in a vacuum concentrator. The dried samples were reconstituted in 20 μL of eluent buffer (0.1% formic acid). Peptide mixtures were analyzed by liquid chromatography tandem mass spectrometry (LC-MS/MS) using an EASY-nLC system (Proxeon Biosystem, Odense, Denmark). Peptides separations were performed on an PicoFrit C18 column (20 cm x 75 μm x 5 μm, New Objective, Massachusetts, USA) for RPLC in positive ionization mode. The mobile phase was composed of: A- water + 0.1% formic acid, and B- acetonitrile + 0.1% formic acid, being made a 2-90% acetonitrile gradient. Injection volumes of 4.5 μL and a flow rate of 300 nL/min over 45 min. Mass spectra were acquired using a LTQ Velos Orbitrap (Thermo Fisher Scientific, Massachusetts, USA) mass spectrometer with ESI ionization operated in positive mode. The following parameters were used to register mass spectra: nanoelectrospray voltage: 2500 V; source temperature: 200 °C; resolution: 60,000; MS range (full scan): 300.0-2,000.0 Da. The mass spectrometer was programmed to perform data acquisition in data-dependent acquisition (DDA) mode. The Orbitrap analyzer was set to generate the spectra after accumulation of a target value of $1e^6$ and the 20 most intense peptide ions with charge states ≥ 2 were sequentially isolated to a target value of 5,000 and fragmented in the linear ion trap by low-energy collision-induced dissociation (CID) (normalized collision energy of 35%). The signal threshold for triggering an MS2 event was set to 1,000 counts. Dynamic exclusion was enabled with an exclusion size list of 500, exclusion duration of 60 s, and repeat count of 1. An activation q of 0.25 and an activation time of 10 ms were used[1].

**MS Data Analysis**

Peak lists were generated from the raw files using Proteome Discoverer version 1.3 (Thermo Fisher Scientific, Massachusetts, USA) with Sequest search engine and searched against the Human International Protein Database (IPI) v. 3.86. Modifications were set as carbamidomethylation (+57.021 Da) as fixed modification, oxidation of methionine (+15.995 Da) as variable modification, one trypsin missed cleavage, and a tolerance of 10 ppm for precursor and 1 Da for fragmented ions. Cross-correlation (XCorr) cutoffs used were +1>1.8, +2>2.2, +3>2.5 and +4>3.5. Using the software Mascot (Matrix Science, London, United Kingdom) and Scaffold Q+ (Proteome Software, Oregon, USA), the MS/MS-based peptide and protein identifications were validated. Peptide and protein identifications were accepted if they had a probability greater than 95% and 99%, respectively, as determined by the Peptide and Protein Prophet algorithm[2].

**Cell Differentiation**

For the differentiation of cardiomyoblasts in cardiomyocytes, FBS concentration was reduced to 1% and retinoic acid 10 nmol L-1 was added to the culture medium (differentiation medium). The medium was changed every 2-3 days. During the process of differentiation, images were acquired for the morphological alteration visualization.

**REFERENCES**

(1) Rubio, M. V.; Zubieta, M. P.; Franco Cairo, J. P.; Calzado, F.; Paes Leme, A. F.; Squina, F. M.; Prade, R. A.; de Lima Damásio, A. R. Mapping N-linked glycosylation of carbohydrate-active enzymes in the secretome of Aspergillus nidulans grown on lignocellulose. *Biotechnol Biofuels* **2016**, *9*, 168. DOI: 10.1186/s13068-016-0580-4.

(2) Ma, K.; Vitek, O.; Nesvizhskii, A. I. A statistical model-building perspective to identification of MS/MS spectra with PeptideProphet. *BMC Bioinformatics* **2012**, *13 Suppl 16* (Suppl 16), S1. DOI: 10.1186/1471-2105-13-S16-S1.
